# Revisiting CPUs for Protein Folding: Xeon-Based Acceleration of AlphaFold2

**DOI:** 10.64898/2026.05.27.728222

**Authors:** Narendra Chaudhary, Wei Yang, Dhiraj Kalamkar, Jianqian Zhou, Soumyadip Ghosh, Lei Xia, Manasi Tiwari, Alexander Heinecke, Bharat Kaul, Sanchit Misra

## Abstract

Protein structure prediction via AlphaFold2 has revolutionized drug discovery, yet its end-to-end execution remains computationally intensive. While GPUs are traditionally favored for deep learning, the AlphaFold2 algorithm consists of heterogeneous phases — preprocessing with sparse database searches and model inference with low-arithmetic-intensity attention modules — that present unique architectural challenges. In this work, we address these bottlenecks by introducing Open-Omics-AlphaFold2, a highly optimized implementation for Intel^®^ Xeon^®^ CPU. By leveraging the CPU’s versatility in handling both sparse preprocessing algorithms and dense matrix operations via Intel Advanced Matrix Extensions (AMX), we accelerate the entire pipeline end-to-end. Our optimization strategy employs multi-level parallelism — spanning multiprocessing, multi-threading, and vectorization — alongside cache-aware tiling and operator fusion. Our results demonstrate that, on a Xeon CPU, Open-Omics-AlphaFold2 achieves 2 7.58 speedup for preprocessing and 19.8 29.2 speedup for model inference over baseline Deepmind-AlphaFold2. Moreover, for a proteome of 391 proteins, Open-Omics-AlphaFold2 running on a dual-socket Intel Xeon 6980P system achieves a remarkable 76% higher throughput over the state-of-the-art GPU-accelerated solution, FastFold, running on a single-socket Intel Xeon 6980P CPU with an NVIDIA H100 offload.

**Code availability:** Baremetal: https://github.com/IntelLabs/open-omics-alphafold Containerized: https://github.com/IntelLabs/Open-Omics-Accelera tion-Framework/tree/main/pipelines/alphafold2-based-protein-folding

## 1 Introduction

Proteins carry out most of the essential functions in biology, and their three-dimensional structures are key to their function. While experimental techniques such as X-ray crystallography [1] and cryo-electron microscopy [2] can determine protein structures with high accuracy, they remain time- and resource-intensive. In comparison, determining a protein’s amino acid sequence is faster and cheaper. Consequently, the task of predicting protein structure directly from sequence, i.e., the protein folding problem [3], has long been considered a central challenge in computational biology.

Over the past five decades, numerous efforts have attempted to solve this problem, but only recently has a breakthrough been achieved with the advent of AlphaFold2 (monomer and multimer) [4, 5]. AlphaFold2 has not only become the de facto standard for protein structure prediction, but has also catalyzed new applications ranging from drug discovery and protein design to the interpretation of genetic variants. AlphaFold2 has also found applications in areas of target prediction, protein function prediction, protein-protein interaction, and biological mechanism of action [6]. Its widespread adoption has led to large-scale deployments across both academia and industry.

Despite its success, AlphaFold2’s end-to-end execution remains computationally demanding. The folding time varies from minutes to hours, depending on protein length, even when run on top-tier GPU-accelerated systems. Additionally, emerging generative AI and agent-based frameworks for drug discovery and protein design often invoke AlphaFold2 multiple times in their pipelines. Therefore, its execution time and cost are expected to become an even bigger bottleneck. Hence, there is a growing need for faster and more cost-efficient solutions for AlphaFold2-based folding.

AlphaFold2 monomer pipeline takes a protein sequence as input and predicts its 3D structure through deep learning-based inference. As illustrated in Figure 1, the pipeline consists of two major stages: preprocessing and model inference. The preprocessing stage, executed on CPUs, performs database searches to find similar protein sequences, followed by multiple sequence alignment (MSA) between the input sequence and the sequences found in the database search. The underlying principle is evolutionary conservation: residues critical to structure and function are likely conserved and thus align well across homologous sequences. This alignment is used to construct the initial “MSA representation”. Simultaneously, structural templates for similar sequences are retrieved to build the initial “Pair representation”, a proxy for inter-residue contacts. The model inference stage employs transformer-based neural networks, the Evoformer and Structure modules, which refine these representations and predict the 3D atomic coordinates. Evoformer applies iterative attention-based and other transformations to both MSA and pair representations, and the Structure module computes the final atomic positions. AlphaFold2-Multimer algorithm – predicts the structure of protein complexes – and follows the same core architecture as the AlphaFold2 algorithm with modules better adapted to protein complexes.

**Figure 1:**
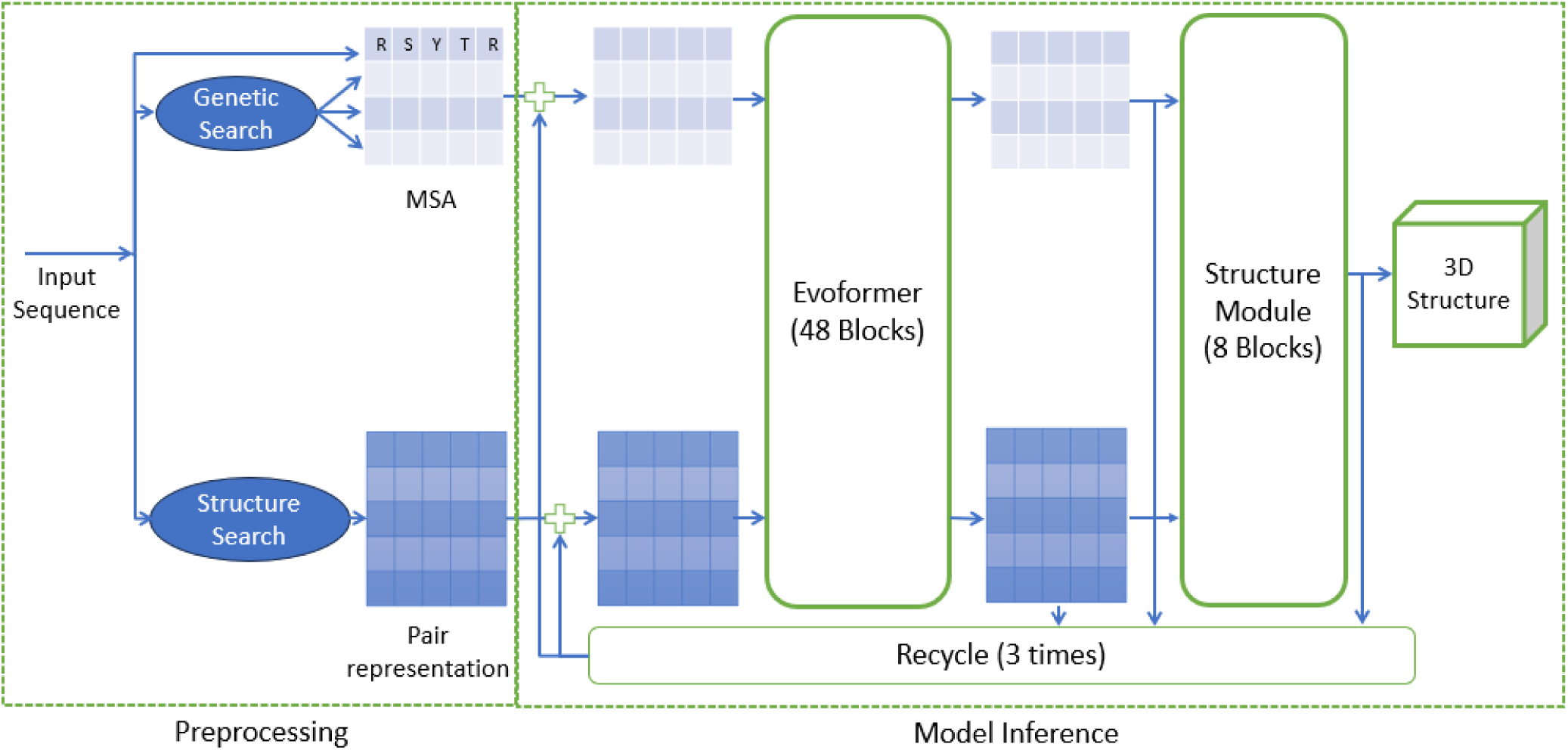
AlphaFold2 protein structure prediction pipeline.

We make the following key observations about AlphaFold2:

(1) **Preprocessing is done on CPUs** and involves file I/O, database searches, and MSA computations – all operations that are known to be better suited to CPUs.
(2) **Preprocessing dominates run time** for AlphaFold2 monomer for shorter proteins (<1000 residues), which represent the majority of naturally occurring proteins [7], while model inference dominates run time for longer proteins (Table 1).
(3) **The attention modules within Evoformer are a performance hotspot** (Figure 3). However, unlike large language models (LLMs), AlphaFold2 uses a relatively small number of attention heads (4 or 8) and small head dimensions (8, 16, 32), which leads to reduced available parallelism in matrix multiplication modules and increased relative time spent on other operations, such as softmax.

**Table 1:**
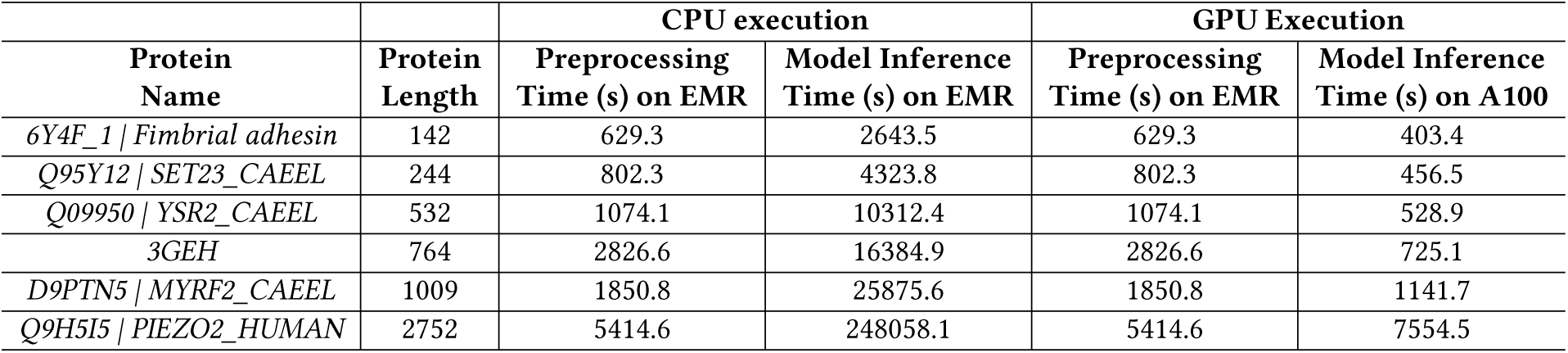
Deepmind-AlphaFold2 performance for single protein inference task on CPU and GPU. We use the default setting of running the inference model 5 times and picking the best out of the 5 predicted structures. Inference time here is sum of the time for the 5 models. Model inference is the primary bottleneck on CPU due to inefficient implementation. With more efficient model inference, as on GPU, preprocessing also becomes a critical bottleneck.

| Protein Name | Protein Length | CPU execution |  | GPU Execution |  |
| --- | --- | --- | --- | --- | --- |
|  |  | Preprocessing Time (s) on EMR | Model Inference Time (s) on EMR | Preprocessing Time (s) on EMR | Model Inference Time (s) on A100 |
| 6Y4F_1 \| *Fimbrial adhesin* | 142 | 629.3 | 2643.5 | 629.3 | 403.4 |
| Q95Y12 \| *SET23_CAEEL* | 244 | 802.3 | 4323.8 | 802.3 | 456.5 |
| Q09950 \| *YSR2_CAEEL* | 532 | 1074.1 | 10312.4 | 1074.1 | 528.9 |
| 3GEH | 764 | 2826.6 | 16384.9 | 2826.6 | 725.1 |
| D9PTN5 \| *MYRF2_CAEEL* | 1009 | 1850.8 | 25875.6 | 1850.8 | 1141.7 |
| Q9H5I5 \| *PIEZO2_HUMAN* | 2752 | 5414.6 | 248058.1 | 5414.6 | 7554.5 |

These insights have surprising architectural implications. While conventional deep learning workloads benefit greatly from GPU-accelerated matrix multiplications (via Tensor Cores), AlphaFold2’s unique attention patterns do not leverage such hardware optimally. Instead, acceleration requires a different system balance: one with strong CPU performance for preprocessing and sufficient – but not necessarily massive – compute for matrix operations.

Modern Intel Xeon CPUs may offer a better balance for such applications. They combine high preprocessing throughput with hardware acceleration for matrix operations via Advanced Matrix Extensions (AMX). This makes them promising candidates for accelerating AlphaFold2 pipeline end-to-end.

Numerous implementations of AlphaFold2 exist today. The original version by DeepMind supports both monomer and multimer prediction [4, 5], using CPUs for preprocessing and GPUs for inference. OpenFold [8], a faithful PyTorch re-implementation, supports CPU or GPU for inference but suffers from inefficient CPU performance. FastFold [9] is a GPU-optimized variant for inference that improves both throughput and memory efficiency, enabling inference on proteins up to 10,000 residues. Work such as [10] has demonstrated AlphaFold2 acceleration on CPU clusters, primarily through resource scheduling and attention re-engineering. However, to our knowledge, no prior work has systematically exploited the architectural features of Intel Xeon CPUs to accelerate AlphaFold2 inference.

In this work, we present Open-Omics-AlphaFold2 – an efficient AlphaFold2 inference pipeline for Intel Xeon CPUs – that achieves state-of-the-art performance without GPUs and without losing accuracy. Our approach combines:

- Mixed-precision execution using BFloat16 and FP32,
- Multi-level parallelism via multiprocessing, multi-threading, and vectorization,
- Hardware acceleration for deep learning with AMX,
- Cache-aware tiling and kernel fusion, and
- A flash-attention-3 inspired implementation of Evoformer attention modules.

Our results demonstrate that, on a Xeon CPU, Open-Omics-AlphaFold2 achieves 2 7.58 speedup for preprocessing and 19.8 29.2 speedup for model inference over baseline Deepmind-AlphaFold2. For instance, for a 2752-length protein, we reduce the preprocessing latency from approximately 90 mins to just under 12 mins and the inference latency from nearly 69 hours to just 2.7 hours. Moreover, for a proteome of 391 proteins, Open-Omics-AlphaFold2 running on a dual-socket Intel Xeon 6980P system achieves a remarkable 76% higher throughput over a hybrid setup comprising one Intel Xeon 6980P CPU with one NVIDIA H100 GPU running our accelerated preprocessing implementation on CPU and FastFold on GPU.

The rest of this paper is organized as follows. In the next section, we provide a background on the AlphaFold2 algorithm. In the subsequent section, we present a high level overview of our acceleration strategies and performance results. This is followed by conclusions, discussion and a detailed explanation of our methods. The paper also has a supplementary section with more details.

## 2 Background on Alphafold2 Algorithm

The AlphaFold2 algorithm consists of the following stages: preprocessing and model inference. Model inference can be further divided into Evoformer and Structure prediction stages. We provide details of the three stages below.

### 2.1 Preprocessing

The goal of the preprocessing stage in AlphaFold2 is to provide the Evoformer module with informative features that reflect which amino acid residues in the input protein sequence are likely to be structurally or functionally important, and to offer initial cues about the protein’s three-dimensional conformation. The key biological insight behind this step is that many proteins across different species share conserved sequences or subsequences due to common ancestry, often linked to similar structure and function. Residues that are conserved across these homologous sequences are likely to play critical structural roles. Moreover, experimentally resolved structures of homologous proteins can serve as valuable templates to guide inference of structure of a protein.

To identify such homologous sequences, AlphaFold2 performs sequence similarity searches across multiple large-scale protein databases using the following tools:

- **JackHMMER**[11] is an iterative profile HMM-based search tool from the HMMER suite. In AlphaFold2, it begins by building a profile HMM from the input query sequence. This profile is used to search sequence databases such as MGnify[12] and UniRef90 [13] for homologous sequences. Matches are used to refine the profile HMM across multiple iterations, enabling the detection of more distant homologs. Each match is aligned to the query via the profile HMM, and these alignments are stacked relative to the query to produce an MSA-like structure.
- **HHblits** [14], part of the HH-suite, further enhances the search process. It constructs a profile HMM from the query sequence and performs **HMM-HMM alignment** against a database of precomputed profile HMMs (e.g., BFD), allowing the identification of remote homologs with greater sensitivity than sequence-HMM methods like JackHMMER. The resulting alignments are again stacked relative to the query to form an MSA-like representation.

The MSA-like outputs from JackHMMER and HHblits are merged to form the final **MSA representation**, which encodes evolutionary co-variation patterns that inform the model’s structure prediction.

For template search, AlphaFold2 uses **HHsearch** [15] – another tool from HH-suite. HHsearch creates a profile HMM of the query and searches through HMMs of sequences with known structures. For the matching sequences, it fetches the corresponding 3D coordinates and uses them to create the **Pair representation**, guiding the prediction process with prior structural knowledge.

In current implementations of AlphaFold2, these tools are applied sequentially. JackHMMER is first used to identify distant homologs, followed by HHblits to expand the MSA and refine the alignment. The resulting MSA information is then combined to generate an **MSA representation**. Subsequently, HHsearch is employed for template searching in structure database, and the identified templates are used to construct a **Pair representation**. Both the **MSA representation** and the **Pair representation** are then passed to the subsequent stage of the inference pipeline – model inference. In the model inference stage, these representations are refined by the AlphaFold2 neural network, which consists of multiple Evoformer blocks, ultimately enabling accurate protein structure prediction.

### 2.2 Evoformer Block

In the AlphaFold2 architecture, the **Evoformer block** (Figure 2) forms the core computational trunk of the network. A typical AlphaFold2 model contains **48 Evoformer blocks** stacked sequentially. Each Evoformer block takes as input the MSA and Pair representations in the form of embeddings. These embeddings are iteratively refined as they pass through the Evoformer stack.

**Figure 2:**
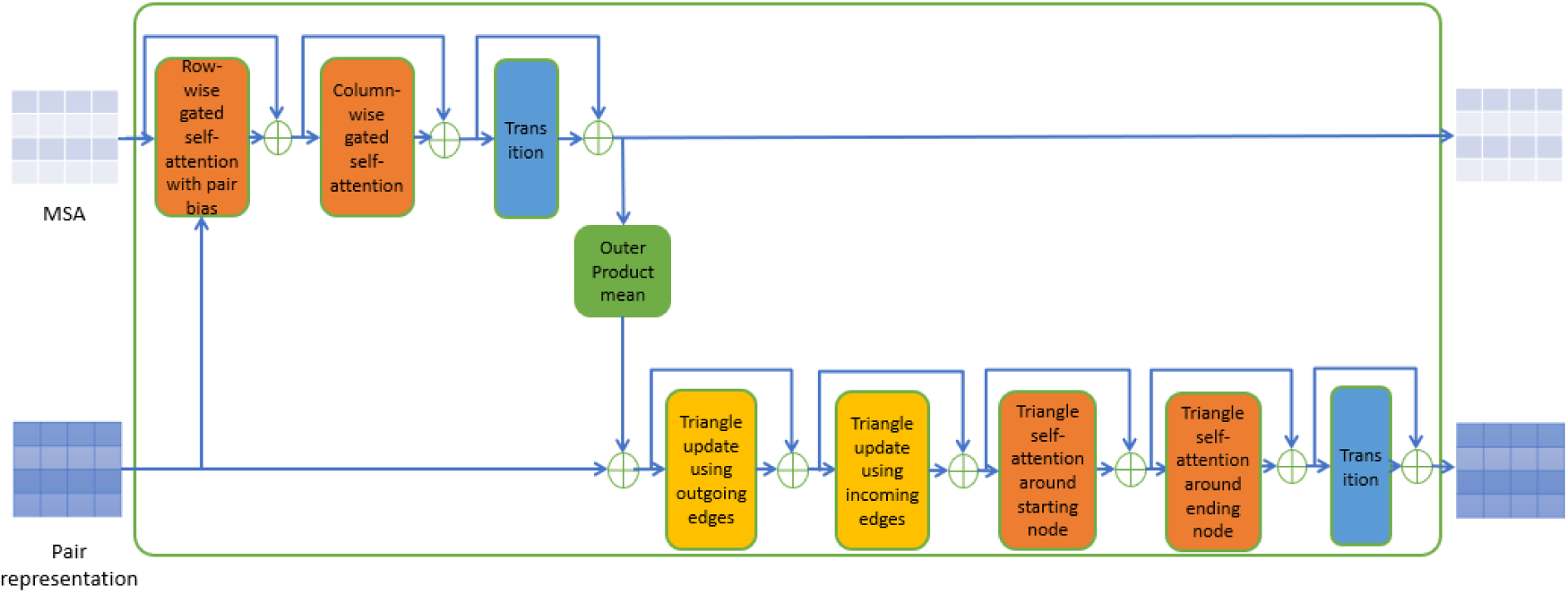
Interior of one Evoformer module containing several sub-modules.

The Evoformer is a *Transformer*-inspired architecture, sharing common elements such as **attention mechanisms** and **multilayer perceptrons (MLPs)**. However, it diverges from the classical Transformer design in several important ways, introducing architectural innovations tailored to protein structure prediction.

*Key Differences from Standard Transformer Blocks:*

(1) **Triangle Modules**: The Evoformer includes specialized *triangle attention* and *triangle update* modules. These are designed to enforce geometric consistency in pairwise residue relationships by modeling *triangle inequality constraints*, which are essential for predicting accurate 3D structures.
(2) **Cross-path Information Flow**: Evoformer maintains two parallel data streams (MSA and pair embeddings) and enables communication between them using operations such as *outer product mean* and *transition layers*.
(3) **2D Criss-Cross Attention**: Attention is applied both *row-wise* and *column-wise* to the MSA and pair embeddings. This strategy, called *2D criss-cross attention*, helps propagate contextual information within sequence and across species.

### 2.3 Structure Module

Structure module is the final stage of the AlphaFold2 inference pipeline. It takes as inputs the pair representation embedding and first row from the MSA embedding tensor. It passes these inputs through a series of specialized modules, most notably the **Invariant Point Attention (IPA)** module, in order to predict the protein structure.

### 2.4 Recycling, multiple models and Computational Load

During inference, the Evoformer and structure modules are executed repeatedly. This is called *recycling*:

- In AlphaFold2 monomer, the Evoformer module is run **192 times** for a single protein (48 blocks × 4 recycling passes).

In AlphaFold2-Multimer, Evoformer module may run up to **960 times**, due to as many as 19 additional recycling passes.

Moreover, AlphaFold2 monomer provides five different models that were trained with slightly different configurations. By default, the system runs the input protein sequence through each of these five models independently and picks the best output structure. AlphaFold2-multimer also provides five different models but, by default, runs each model five times with different seeds. Therefore, it generates 25 candidate structures and picks the best from them. Therefore, multimeric protein inputs result in more recycles and also more model executions to produce a larger number of candidates. Moreover, multimeric proteins consist of multiple chains, resulting in longer total sequence lengths within the Evoformer, which increases computational demands. As a result, model inference constitutes a much larger fraction of the overall structure prediction time in multimer cases.

## 3 Results

In this section, we present the results of our experiments. We begin by reporting the baseline performance of Deepmind-AlphaFold2 (original JAX-based implementation) on both CPU and GPU platforms. Next, we reproduce Deepmind-AlphaFold2 code in PyTorch as PyTorch-Baseline-AlphaFold2 and we use it as a starting point for our optimizations. We conduct a performance characterization of AlphaFold2, identifying key computational bottlenecks and providing an overview of our proposed acceleration strategy. Subsequently, we present the performance gains achieved, focusing on critical components including preprocessing, gating attention, and triangle multiplication kernels. We first provide results for accelerated folding of individual proteins by comparing Deepmind-AlphaFold2 and Open-Omics-AlphaFold2. Finally, we present the results for folding of multiple proteins and compare the performance of Open-Omics-AlphaFold2 (optimized CPU implementation), against FastFold [9], a state-of-the-art optimized GPU-based version, highlighting the effectiveness and efficiency of our approach.

### 3.1 Experimental Setup

#### 3.1.1 Software

For all our experiments, we follow the same software configuration and settings as described in the AlphaFold2 paper [4]. During preprocessing, we use JackHMMER [11], HHblits [14] and HHsearch with identical settings to those used in Deepmind-AlphaFold2. We also use the official model parameters from AlphaFold2 and AlphaFold2-Multimer (version v3) for all inference tasks. Detailed version information for the software, tools, and codebases used in our experiments is provided in Table S1.

#### 3.1.2 Protein database and sequence datasets

The AlphaFold2 pipeline relies on several genetic and structure databases, including BFD, MGnify, UniRef30, UniRef90, PDB70, and the PDB. For AlphaFold2-Multimer, two additional databases are required: PDB seqres and UniProt. Collectively, these databases occupy approximately 3 terabytes of storage. We evaluate performance using the same database snapshot across all AlphaFold2 implementations to ensure consistency and fair comparison. List of these genetic databases and their download link is provided in Table S2.

Protein inference execution time and memory consumption depends heavily on the sequence length of a protein. Therefore, to assess performance for AlphaFold2 monomer with respect to the protein length, we present results of Deepmind-AlphaFold2, Fast-Fold, PyTorch-Baseline-AlphaFold2, Open-Omics-AlphaFold2 on individual real proteins of different lengths {108, 244, 532, 764, 1009, 2752}. We believe this set of protein lengths is representative of the range of protein lengths in nature. To also demonstrate the performance of Open-Omics-AlphaFold2 on a set of proteins as opposed to single proteins, we compare it with FastFold on a subset of 391 real proteins from *C. elegans* proteome. The protein lengths in this subset range from 31 to 1001 amino acid residues reflecting the lengths of majority of proteins found in nature. For AlphaFold2-multimer, we use four individual protein complexes of lengths {530, 716, 1252, 2216} and set of 6 real protein complexes. We download the *C. elegans* proteome [16] from Uniprot webpage [17], sort the list of proteins by protein length and select every 10th protein from the sorted list. We only select protein up to the length of 1001 residues. Some proteins failed during preprocessing with the original source code and we therefore remove them from the list. We downloaded other proteins from the protein Data Bank [18].

#### 3.1.3 Hardware

Our experimental evaluation utilizes two generations of Intel^®^ Xeon^®^ CPUs and NVIDIA GPUs. Specifically, we employ the Intel^®^ Xeon^®^ Platinum 8592+ (Emerald Rapids, **EMR**) and the Intel^®^ Xeon^®^ 6980P (Granite Rapids, **GNR**), alongside NVIDIA **A100** and **H100** GPUs. The EMR, GNR, and H100 benchmarks were conducted on on-premises systems, while A100 experiments utilized Google Cloud Platform (GCP) a2-highgpu-1g instances. To mitigate I/O bottlenecks during preprocessing, all experiments were conducted using Solid State Drives (SSDs). Comprehensive system specifications are detailed in Tables S3 and S4.

Both EMR and GNR are dual-socket configurations featuring a total of 128 and 256 physical cores, respectively. The EMR system is equipped with 1 TB of DDR5 memory (717 GB/s peak bandwidth), while the GNR system utilizes 1.5 TB of MRDIMM memory, with a peak bandwidth of 1690 GB/s. Each core integrates two 512-bit SIMD units (AVX512). While these units support general-purpose vectorization, both architectures include **Intel® Advanced Matrix Extensions (AMX)**—dedicated hardware blocks designed to accelerate low-precision matrix multiplication. AMX significantly outperforms standard SIMD units for deep learning kernels, providing peak Bfloat16 throughput of 262 TFLOPS on EMR and 524 TFLOPS on GNR. Matrix multiplication with FP32 precision needs to use AVX512 units while with Bfloat16 precision, it can also use the AMX units.

The NVIDIA A100 and H100 GPUs feature 40 GB of HBM2 and 80 GB of HBM3 memory, with peak bandwidths of 1555 GB/s and 3350 GB/s, respectively. These units leverage **Tensor Cores** for specialized matrix acceleration. The A100 delivers 156 TFLOPS (TF32) and 312 TFLOPS (Bfloat16) of peak performance. The H100 increases these capabilities to 495 TFLOPS (TF32) and 990 TFLOPS (Bfloat16)^1^.

### 3.2 Performance Characterization of Baseline AlphaFold2

To guide our acceleration efforts, we first run Deepmind-AlphaFold2 on both CPU and GPU. Table 1 shows baseline performance on both platforms. CPU runs were performed entirely on EMR, while GPU runs used EMR for preprocessing and A100 for inference.

On CPU, model inference dominates runtime and is significantly slower than on GPU due to inefficiencies in the implementation. The original JAX-based Deepmind-AlphaFold2 is especially inefficient on CPUs at longer protein lengths. Therefore, we first port AlphaFold2 codebase to PyTorch to establish a better CPU baseline. We call this new CPU baseline as PyTorch-Baseline-AlphaFold2. Next, we focus on accelerating CPU inference from this baseline. On GPU, preprocessing accounts for 40–80% of total time and is a major bottleneck. Once inference is optimized on CPU, preprocessing is expected to become the next bottleneck, motivating us to accelerate it as well.

To further analyze model inference, we profiled the runtime of individual components for a representative protein of length 764 (Figure 3). The Evoformer blocks –specifically the “Extra MSA Evoformer” and “No Extra MSA Evoformer” – together account for nearly 90% of the total inference time. These blocks consist of attention-based sub-modules (row-wise and column-wise attention, triangle attention) and other modules such as transition, outer product mean, and triangle update.

**Figure 3:**
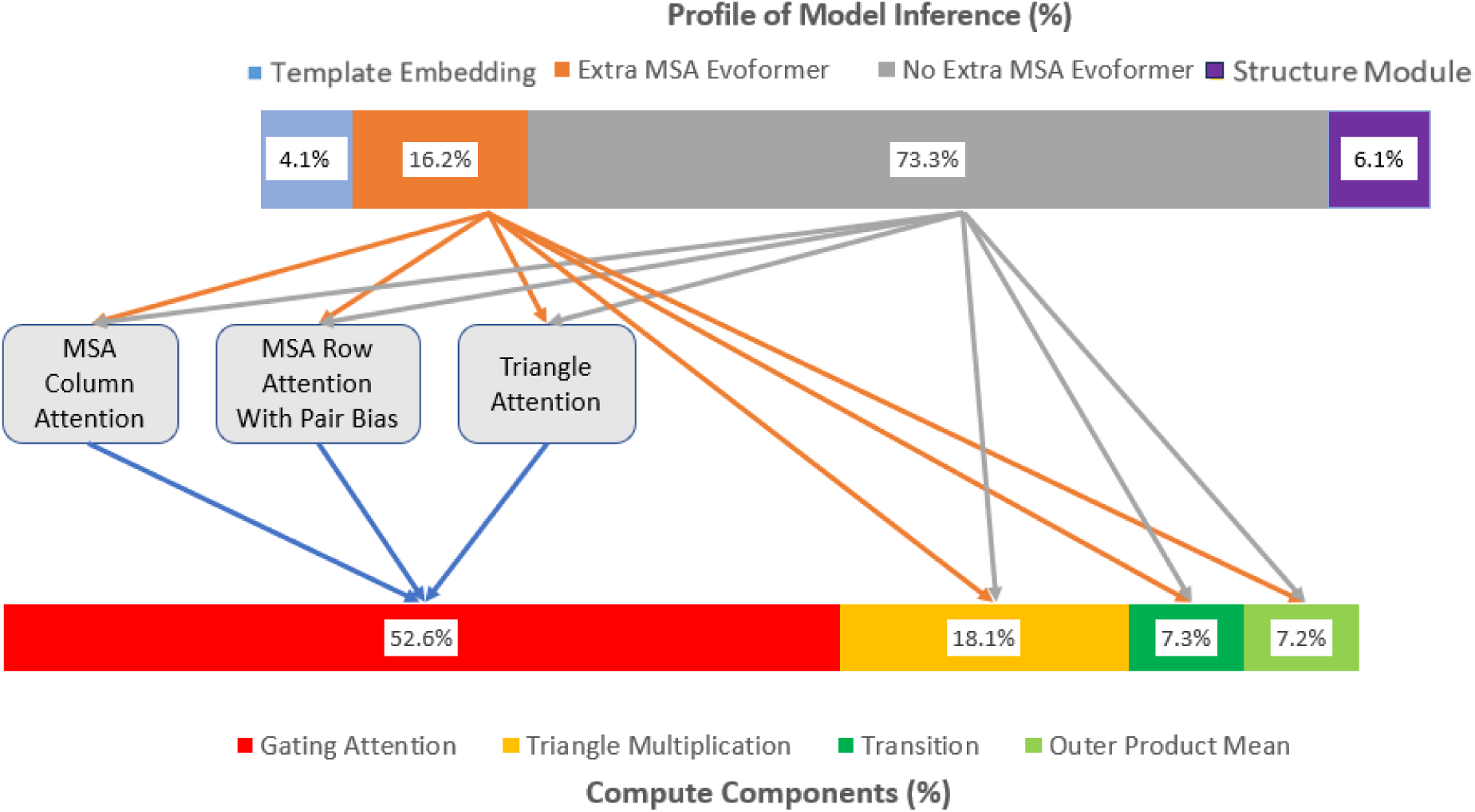
Runtime breakdown of PyTorch-Baseline-AlphaFold2 model inference for a protein of length 764. The “Extra MSA Evoformer” and “No Extra MSA Evoformer” dominate the total execution time, with the gating attention and triangle multiplication sub-modules together accounting for 70% of total model inference time.

All attention-based sub-modules in the Evoformer block are implemented using a single generalized *Gating Attention* module (see Algorithm 1), with variations in the number of heads and head dimensions. Similarly, all triangle update modules are implemented using a single *Triangle Multiplication* module (see Algorithm 2). The gating attention and triangle multiplication sub-modules together dominate the runtime, contributing approximately 70% of the total inference time for the selected protein. This is largely due to their computational complexity: multi-head attention scales as *S*^2^ [19–21], where *S* is the sequence length. In many attention layers of AlphaFold2, the batch dimension is same as the sequence length, effectively making the complexity of these layers *S*^3^. The triangle multiplication module also scales as *S*^3^. Unoptimized implementations of attention module have a space complexity O(*S*^2^) in sequence length. Therefore, memory consumption can quickly increase with protein sequence length (Table 5). Although profiles will vary with protein length, these two sub-modules are expected to remain the primary bottlenecks becoming even more prominent for longer sequences and multimer proteins due to their cubic scaling.

### 3.3 A Brief Overview of Our Acceleration

Broadly, we apply the following approaches to accelerate the pipeline (illustrated in Figure 4):

**Figure 4:**
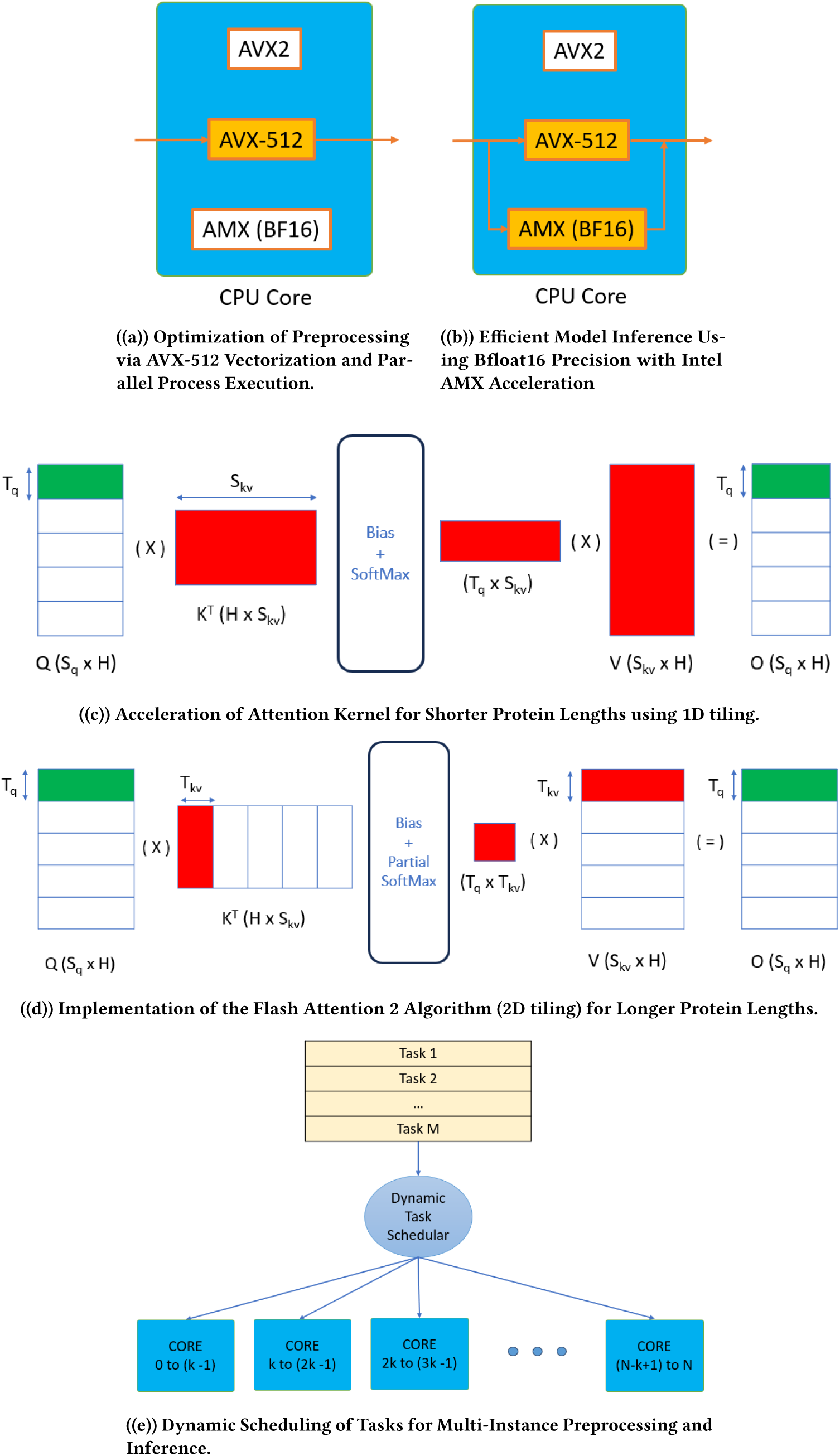
Illustration of Key Acceleration Techniques Employed in This Work.

#### 3.3.1 Multi-processing with dynamic scheduling

Folding of a single protein does not scale linearly with the number of CPU cores for several reasons. First, many proteins are too small to generate sufficient computational work to keep all cores occupied. Second, preprocessing involves searches over large databases, during which, cores frequently idle while awaiting disk access. Third, the software tools used for preprocessing scale poorly with increasing core counts. Finally, unavoidable sequential steps further constrain multi-core efficiency. Consequently, modern CPUs with a few hundred cores often remain underutilized.

At the same time, the ample memory available on these processors and our optimizations to reduce memory consumption discussed in Section 5.5.2, enables the simultaneous analysis of many proteins. Because the folding of each protein is independent, this problem is *embarrassingly parallel*. Therefore, we fold multiple proteins concurrently, dividing the available cores on a CPU equally among them. This strategy is particularly effective in proteome-scale analyses and protein design, where many structures may need to be determined.

Execution time varies markedly across proteins: inference scales cubically with protein length, whereas preprocessing time depends on the number of database matches. As a result, there is significant load imbalance across proteins in both preprocessing and inference and it is especially pronounced during inference. If not handled well, it may lead to certain cores finishing early and certain cores taking too long to finish. To address this, we use two-pronged approach. First, we separate folding into two phases – (i) preprocessing of all proteins, with intermediate outputs written to disk, and (ii) model inference – due to their vastly different load imbalance characteristics. Second, within each phase, we adopt dynamic scheduling. Multiple proteins are processed simultaneously, and once a protein completes, its cores are immediately assigned to a new one. Further implementation details are provided in Supplementary Section S3.1. Model inference on multiple proteins simultaneously became possible only due to the application of our optimizations that reduce the memory footprint for a single protein. The vanilla PyTorch/JAX code requires a large amount of RAM, which would otherwise prevent the launch of multiple processes on a CPU.

#### 3.3.2 Acceleration of Preprocessing

We accelerated preprocessing by fine-tuning the performance of HHblits, JackHMMER and HHsearch through a combination of 1) moving from 128/256-bit vectorization to 512-bit vectorization of hotspots where applicable, 2) using the appropriate compiler flags to generate an optimized binary, 3) Using the optimal execution setup; e.g. thread-affinity settings. Additionally, baseline AlphaFold2 runs these tools sequentially. Given that the individual tools do not scale well with large number of cores, we run them in parallel using a subset of available cores for each. Supplementary Section S3.2 provides more details. In future work, we plan to do further optimizations of the preprocessing tools.

#### 3.3.3 Acceleration of Model Inference

We primarily focus on accelerating inference since that is the most inefficient stage on CPU. As mentioned in Section 3.2, the majority of inference time is spent in two generalized modules – *Gating Attention* and *Triangle Multiplication*. We accelerate these modules through a combination of Flash attention 2 inspired cache optimizations, operator fusion and use of reduced precision (BFloat16 instead of FP32). Switch to BFloat16 also enables us to use the hardware support for deep learning named AMX [3.1.3] that provides up to 16 higher performance for matrix multiplication compared to FP32 precision that needs to use AVX512-based vectorization for matrix multiplication. Section 5 provides more details.

### 3.4 Performance Evaluation

#### 3.4.1 Preprocessing

The performance improvements for individual tools and the end-to-end preprocessing stage are summarized in Table 2. In preprocessing, majority of time is spent on HHblits and JackHMMER with HHsearch consuming a tiny portion of total time. Therefore, we skip reporting the speedups obtained on HHsearch individually. While we observe substantial gains in individual components – up to 2× for JackHMMER and 10× for HHblits – the most significant impact is seen at the level of total preprocessing time. Notably, our optimizations provide higher performance gains for longer proteins: while shortest sequence sees a 2× improvement, the speedup scales remarkably to **7.58** for the longest sequence (PIEZO2_HUMAN, 2752 residues). This scaling behavior shows that our transition to 512-bit vectorization and parallel tool execution effectively mitigates the sequential bottlenecks that traditionally dominate preprocessing for longer proteins.

**Table 2:**
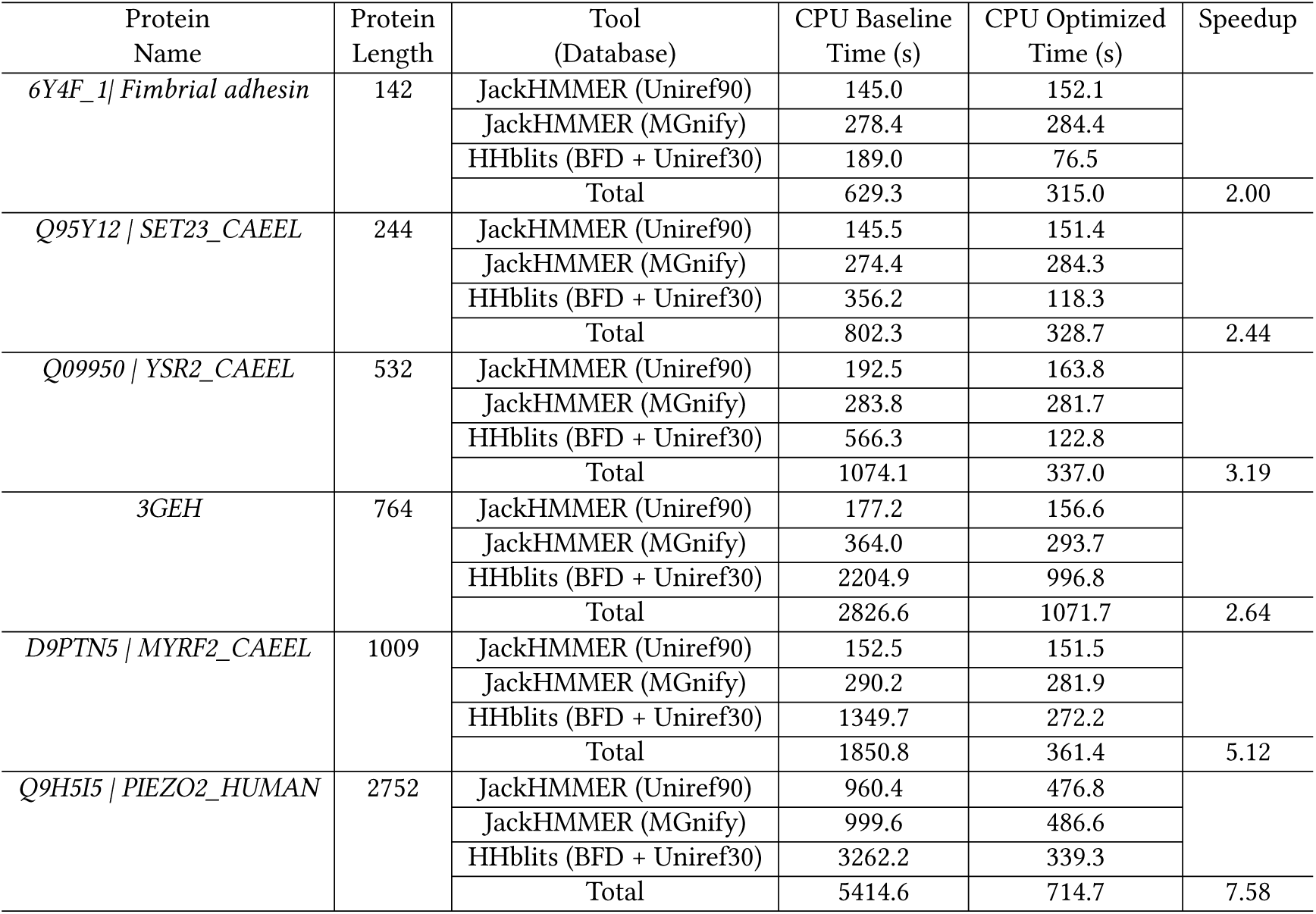
Performance improvements in the AlphaFold2 preprocessing stage. "Total" in the CPU Baseline column presents results for a baseline run in which individual tools are run sequentially, while "Total" in the CPU Optimized column shows the end-to-end time with parallel execution of tools. We ran these experiments on a single-socket EMR CPU machine.

#### 3.4.2 Inference Kernels

This section details the performance improvements achieved for the *Gating Attention* and *Triangle Multiplication* modules on single-socket of EMR CPU that was detailed in Section 3.1.3. The execution time of the attention module is influenced by batch size (B), sequence length (S), number of heads (N), and head size (H). Similarly, the execution time of the triangle multiplication module depends on sequence length (S), embedding dimension (D), and equation type. To evaluate our optimizations, we performed ablation studies with parameter sets commonly used within the Evoformer, for protein lengths of 244, 764, and 2752 residues. For these ablation studies, we created a single-layer neural network with either a gating attention layer or a triangle multiplication layer and measured the time for its forward pass. We disabled gradient computation so that we are measuring only the compute required for the forward pass.

Tables 3 and 4 present the results of performance improvements achieved through various optimization techniques. The “Baseline time” refers to the out-of-the-box run of PyTorch code, that uses the off-the-shelf CPU-optimized oneDNN library underneath, on a single-socket Xeon without any AlphaFold2-specific optimizations. This PyTorch baseline is already more performant than what most users experience with the original Jax based AlphaFold2. In “Improved Config time” column, we apply some improved configuration changes to improve performance. These included the use of torch.compile(), alongside adjustments to environment variables controlling jemalloc, OpenMP thread count, and KMP affinity. We selected the configuration that provided the best results and present those findings in the tables. Finally, “Optimized Time” column represents our accelerated implementation where we replaced standard layers with custom layers implemented as accelerated kernels in C++ as PyTorch extensions.

**Table 3:**
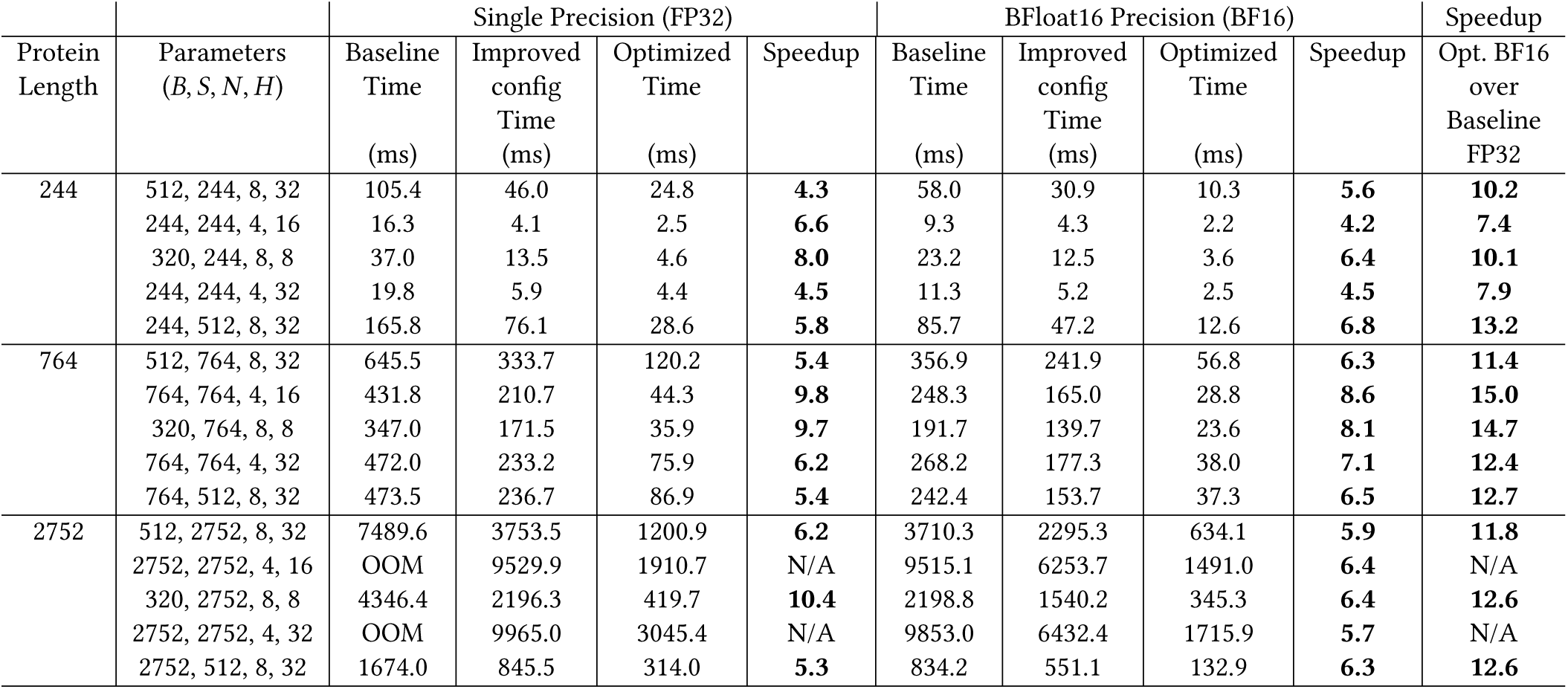
Performance improvements in the gating attention kernel achieved through various optimizations. Speedup is measured relative to the baseline, separately for FP32 and Bfloat16 precision. The last column reports the total speedup achieved: most optimal Bfloat16 implementation vs. baseline FP32 implementation.

| Protein Length | Parameters<br>($B, S, N, H$) | Single Precision (FP32) | | | | BFloat16 Precision (BF16) | | | | Speedup<br>Opt. BF16<br>over<br>Baseline<br>FP32 |
| --- | --- | --- | --- | --- | --- | --- | --- | --- | --- | --- |
|  |  | Baseline Time<br>(ms) | Improved config Time<br>(ms) | Optimized Time<br>(ms) | Speedup | Baseline Time<br>(ms) | Improved config Time<br>(ms) | Optimized Time<br>(ms) | Speedup |  |
| 244 | 512, 244, 8, 32 | 105.4 | 46.0 | 24.8 | **4.3** | 58.0 | 30.9 | 10.3 | **5.6** | **10.2** |
|  | 244, 244, 4, 16 | 16.3 | 4.1 | 2.5 | **6.6** | 9.3 | 4.3 | 2.2 | **4.2** | **7.4** |
|  | 320, 244, 8, 8 | 37.0 | 13.5 | 4.6 | **8.0** | 23.2 | 12.5 | 3.6 | **6.4** | **10.1** |
|  | 244, 244, 4, 32 | 19.8 | 5.9 | 4.4 | **4.5** | 11.3 | 5.2 | 2.5 | **4.5** | **7.9** |
|  | 244, 512, 8, 32 | 165.8 | 76.1 | 28.6 | **5.8** | 85.7 | 47.2 | 12.6 | **6.8** | **13.2** |
| 764 | 512, 764, 8, 32 | 645.5 | 333.7 | 120.2 | **5.4** | 356.9 | 241.9 | 56.8 | **6.3** | **11.4** |
|  | 764, 764, 4, 16 | 431.8 | 210.7 | 44.3 | **9.8** | 248.3 | 165.0 | 28.8 | **8.6** | **15.0** |
|  | 320, 764, 8, 8 | 347.0 | 171.5 | 35.9 | **9.7** | 191.7 | 139.7 | 23.6 | **8.1** | **14.7** |
|  | 764, 764, 4, 32 | 472.0 | 233.2 | 75.9 | **6.2** | 268.2 | 177.3 | 38.0 | **7.1** | **12.4** |
|  | 764, 512, 8, 32 | 473.5 | 236.7 | 86.9 | **5.4** | 242.4 | 153.7 | 37.3 | **6.5** | **12.7** |
| 2752 | 512, 2752, 8, 32 | 7489.6 | 3753.5 | 1200.9 | **6.2** | 3710.3 | 2295.3 | 634.1 | **5.9** | **11.8** |
|  | 2752, 2752, 4, 16 | OOM | 9529.9 | 1910.7 | N/A | 9515.1 | 6253.7 | 1491.0 | **6.4** | N/A |
|  | 320, 2752, 8, 8 | 4346.4 | 2196.3 | 419.7 | **10.4** | 2198.8 | 1540.2 | 345.3 | **6.4** | **12.6** |
|  | 2752, 2752, 4, 32 | OOM | 9965.0 | 3045.4 | N/A | 9853.0 | 6432.4 | 1715.9 | **5.7** | N/A |
|  | 2752, 512, 8, 32 | 1674.0 | 845.5 | 314.0 | **5.3** | 834.2 | 551.1 | 132.9 | **6.3** | **12.6** |

**Table 4:**
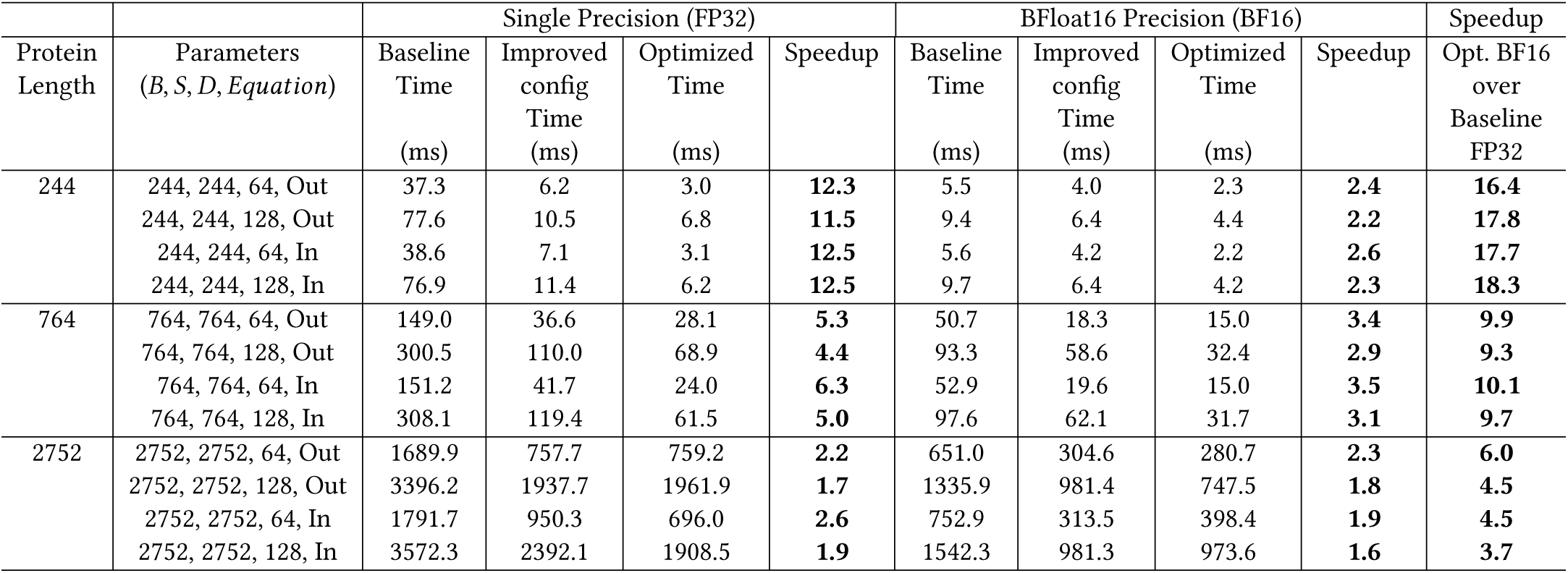
Performance improvements in triangle multiplication kernel through various performance optimizations. Speedup is calculated over baseline separately for FP32 and Bfloat16. The last column reports the total speedup achieved: most optimal Bfloat16 implementation vs. baseline FP32 implementation.

**Table 5:** Memory reduction improvements in largest kernels through tiling optimizations.

| Kernel<br>(Parameters) | Single Precision (FP32) |  |  | BFloat16 Precision (BF16) |  |  | Reduction |
| --- | --- | --- | --- | --- | --- | --- | --- |
|  | Baseline<br>Memory (GB) | Optimized<br>Memory (GB) | Reduction<br>(%) | Baseline<br>Memory (GB) | Optimized<br>Memory (GB) | Reduction<br>(%) | Opt. BF16 vs.<br>baseline FP32<br>(%) |
| Gating Attention<br>(2752, 2752, 4, 32) | 661.9 | 33.7 | 94.9 % | 329.2 | 19.4 | 94.1 % | 97.1 % |
| Triangle Multiplication<br>(2752, 2752, 128, In) | 91.8 | 26.2 | 71.5 % | 45.8 | 15.6 | 65.9 % | 83 % |

Our results demonstrate that basic configuration improvements can provide significant performance gains, but hand-written kernels offer a more significant improvement. Furthermore, utilizing BFloat16 precision can result in a 1.1 2.4 speedup over single precision (Float32) due to AMX. Although AMX can theoretically deliver 16× higher FLOPs with BFloat16 precision over AVX512, that is needed for FP32 precision, if the entire compute is only matrix multiplication, its actual performance is reduced by memory bandwidth limitations and the fact that matrix multiplication is a fraction of the total compute. Overall, we obtained speedups ranging from 3.7 18.3 for the two modules when moving from the baseline FP32 implementations to the optimized Bfloat16 implementations.

Furthermore, our implementation of a flash-attention-based algorithm, along with other optimizations, significantly reduces memory consumption in the gating attention and triangle multiplication modules. Table 5 summarizes the memory savings achieved by these optimized kernels. For the longest protein sequences, we achieve reduction in memory usage of over 97% in gating attention and of 83% in triangle multiplication. These improvements make it feasible to perform long protein structure inference, which was previously challenging due to memory constraints. Moreover, on systems with large memory capacity, such as EMR with 1 TB of RAM, it becomes possible to run multiple concurrent inference jobs for smaller proteins. Specifically, we can efficiently launch more than 16 parallel inference instances on a single machine.

#### 3.4.3 Monomer Inference

The comparison of end-to-end model inference performance of Deepmind-AlphaFold2, PyTorch-Baseline-AlphaFold2 and Open-Omics-AlphaFold2 (integrated with our optimized modules for gating attention and triangle multiplication) on a single socket of EMR is presented in Table 6. Notably, our implementation overcomes the severe scalability limitations of the original JAX-based DeepMind-AlphaFold2 on CPU architectures, achieving speedups of 19.8 29.2 for proteins of lengths 142−2752. In particular, for the 2752-residue protein (PIEZO2_HUMAN), the original implementation required approximately 69 hours (248,058 s) for inference. In contrast, Open-Omics-AlphaFold2 reduces this execution time to roughly 2.7 hours (9,658 s), representing a 25.7× speedup. Furthermore, while the vanilla PyTorch baseline encountered out-of-memory (OOM) errors at this sequence length, our memory-efficient kernels enable seamless execution, effectively democratizing long-protein inference on standard CPU-based workstations.

**Table 6:** Model Inference performance of AlphaFold2 monomer for inference of a single protein on a single EMR socket.

| Protein<br>Name | Protein<br>Length | Deepmind<br>AF2<br>Time (s) | Single Precision (FP32) |  | BFloat16 Precision (BF16) |  | Overall<br>Speedup |
| --- | --- | --- | --- | --- | --- | --- | --- |
|  |  |  | PyTorch<br>Baseline AF2<br>Time (s) | Open-Omics<br>AF2<br>Time (s) | PyTorch<br>Baseline AF2<br>Time (s) | Open-Omics<br>AF2<br>Time (s) |  |
| *6Y4F_1 \| Fimbrial adhesin* | 142 | 2643.5 | 304.8 | 241.9 | 180.5 | 133.5 | 19.8 |
| *Q95Y12 \| SET23_CAEEL* | 244 | 4323.8 | 581.5 | 243.8 | 295.7 | 166.9 | 25.9 |
| *Q09950 \| YSR2_CAEEL* | 532 | 10312.4 | 1479.1 | 609.5 | 949.4 | 353.4 | 29.2 |
| *3GEH* | 764 | 16384.9 | 2789.4 | 1091.7 | 1969.3 | 621.7 | 26.4 |
| *D9PTN5 \| MYRF2_CAEEL* | 1009 | 25875.6 | 4994.0 | 1887.0 | 3734.0 | 1065.3 | 24.3 |
| *Q9H5I5 \| PIEZO2_HUMAN* | 2752 | 248058.1 | OOM | 17618.0 | OOM | 9658.0 | 25.7 |

In addition to performance, we also compare the prediction accuracy of Open-Omics-AlphaFold2 with Deepmind-AlphaFold2. We compare the predicted 3D structures using structural visualization software. Furthermore, Table 7 presents a comparison of pLDDT (predicted Local Distance Difference Test) scores for multiple proteins for Deepmind-AlphaFold2 on CPU and GPU and Open-Omics-AlphaFold2. The results show that the pLDDT scores from Open-Omics-AlphaFold2 are either close to or identical to those from Deepmind-AlphaFold2 on both platforms. Minor differences in pLDDT values can be attributed to slight variations in numerical precision arising from differences in hardware and implementation changes. This experiment further demonstrates that BFloat16 achieves accuracy comparable to FP32, while delivering substantial speedups.

**Table 7:**
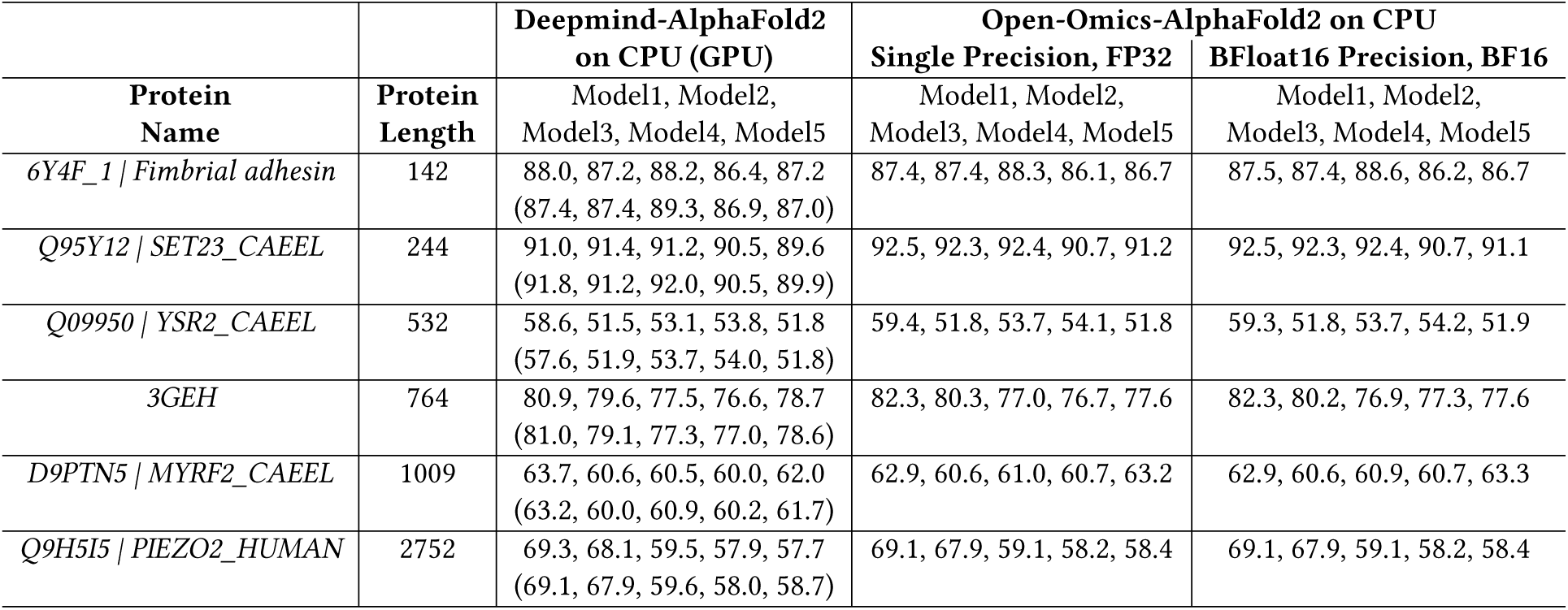
Model Inference mean of plddts values.

#### 3.4.4 Multimer Inference

We have also implemented the Multimer version of the AlphaFold2 model inference in Open-Omics-AlphaFold2, leveraging the optimized kernels. Specifically, we implemented a different version of fused triangle multiplication kernel tailored for AlphaFold2-Multimer.

Table 8 compares the inference times of Deepmind-AlphaFold2 and Open-Omics-AlphaFold2 across a set of multimeric protein complexes. Deepmind-AlphaFold2 was executed on an A100 GPU, while Open-Omics-AlphaFold2 ran on a single-socket CPU (EMR). For each protein, we used the default policy that performs inference across five trained models with five predictions per model, totaling 25 predictions. We only present results for GPU execution for Deepmind-AlphaFold2 due to the prohibitively long runtimes that it would require on CPUs. Results in table 8 highlights the efficiency gains enabled by our CPU-optimized Open-Omics-AlphaFold2.

**Table 8:** Model inference performance of AlphaFold2-Multimer on multimer protein complexes. Deepmind-AlphaFold2 was run on a single A100 GPU, while Open-Omics-AlphaFold2 was executed on a single socket of the EMR CPU platform.

| Protein Name | Protein Length | Multimer Chains | Deepmind-AlphaFold2 GPU Time (s) | Open-Omics-AlphaFold2 CPU Time Bfloat16 (s) | Overall Speedup |
| --- | --- | --- | --- | --- | --- |
| 7KHD \| Tumor necrosis factor | 530 | 4 | 1996.6 | 3032.7 | 0.66 |
| 6IWD \| Human papillomavirus | 716 | 4 | 1656.2 | 2286.4 | 0.72 |
| 6E3K \| Interferon gamma | 1252 | 6 | 6826.3 | 14621.0 | 0.47 |
| 7S81 \| Poly [ADP-ribose] polymerase | 2216 | 16 | 68928.7 | 171719.6 | 0.40 |

#### 3.4.5 Multi-protein Inference

For our final evaluation, we aim to maximize computational throughput and CPU utilization by executing the end-to-end AlphaFold2 pipeline (both preprocessing and model inference) for multiple proteins concurrently via dynamic scheduling.

We benchmark Open-Omics-AlphaFold2 against FastFold, the fastest state-of-the-art GPU-based implementation for end-to-end protein folding. Crucially, because even GPU-centric AlphaFold2 workflows execute the data-preprocessing phase on the host CPU, we isolated this stage during our initial testing. After verifying that the optimized preprocessing pipeline in Open-Omics-AlphaFold2 significantly outperforms the baseline implementation in FastFold, we integrated Open-Omics-AlphaFold2’s preprocessing into the FastFold pipeline as well to ensure an equitable comparison. Furthermore, since FastFold lacks a native BFloat16 implementation, we executed its FP32 pipeline by leveraging hardware-accelerated TF32 on the GPU. While we attempted to transition the FastFold baseline to BFloat16 via automatic mixed precision (autocast), the model suffered from numerical instability and accuracy degradation. Although a BFloat16 implementation of FastFold is fundamentally feasible given careful, manual precision management, it is beyond the scope of this work.

Consequently, we evaluate and compare the following three hardware and software configurations:

(1) **Open-Omics-AlphaFold2 (FP32):** with preprocessing and FP32 model inference executed on a dual-socket GNR platform.
(2) **Open-Omics-AlphaFold2 (BFloat16):** with preprocessing and AMX-accelerated BFloat16 model inference executed on a dual-socket GNR platform.
(3) **FastFold Baseline (TF32):** Open-Omics-AlphaFold2 pre-processing executed on a single-socket GNR platform, combined with FastFold inference executed on an NVIDIA H100 GPU utilizing TF32 precision.

This experimental design ensures that all three setups utilize an equivalent number of physical hardware devices, while granting the GPU baseline both our advanced preprocessing optimizations and its highest-performing available numerical precision (TF32).

As detailed in Section 3.3.1, the preprocessing and model inference phases were launched independently to optimize runtime resource allocation. During the inference phase, we enabled hyper-threading on the CPU, which yielded measurable throughput improvements. GPU-based inference instances for different proteins had to be executed sequentially because FastFold does not have the functionality to run multiple proteins in parallel. Moreover, due to spatial limitations imposed by the H100’s 80 GB memory capacity, it is not possible to run inference on many independent proteins simultaneously.

For the monomer multi-protein experiments, we evaluated structure prediction throughput using a dataset of 391 proteins from the *C. elegans* proteome. For both the preprocessing and inference stages, we launched 24 parallel instances to process 24 proteins concurrently, allocating 10 physical CPU cores per instance. Model inference followed the default AlphaFold2 protocol, executing 5 models with one prediction per model. The throughput results are summarized in Figure 5(a). Remarkably, Open-Omics-AlphaFold2 running in BFloat16 precision across a dual-socket GNR CPU achieves a 1.76 **speedup (a 76% performance gain)** over the hybrid baseline that pairs a single GNR socket for preprocessing with an H100 GPU for inference. Notably, Open-Omics-AlphaFold2 not only performs better in preprocessing, for which it gets double the number of CPU sockets, BFloat16 based inference on GNR also beats TF32 based inference on H100 by nearly 1.6.

**Figure 5:**
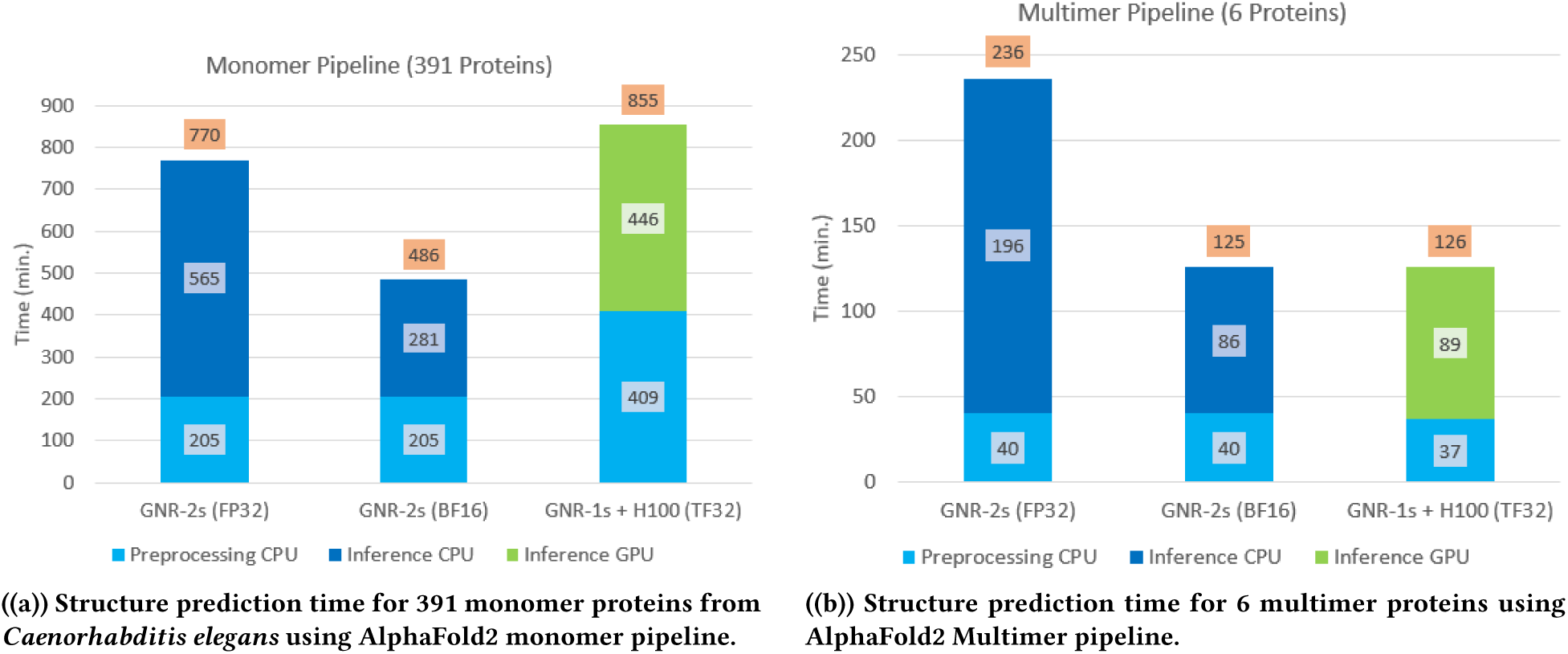
Comparison of execution time for end-to-end AlphaFold2 pipeline for different CPU-GPU systems and precision. GNR-2s (FP32): AlphaFold2 running with FP32 precision with two sockets of GNR. GNR-2s (Bfloat16): AlphaFold2 running with BFloat16 precision with two sockets of GNR. GNR-1s + H100 (TF32): Preprocessing running on one socket of GNR and inference running with TF32 precision on one H100 GPU.

For the multimer multi-protein experiments, we performed evaluations on a dataset comprising 6 distinct protein complexes. During the preprocessing phase, we launched 6 concurrent instances – one per complex – to run all of them in parallel. During the subsequent model inference phase, each complex requires 25 individual inference passes (5 models with 5 predictions per model), yielding a workload pool of 150 total inference tasks. To maximize hardware occupancy, we executed up to 24 inference tasks concurrently. As demonstrated in Figure 5(b), the dual-socket GNR CPU configuration performs competitively with the hybrid single-socket GNR and H100 GPU configuration, highlighting the effectiveness of our CPU-targeted co-design under dense, multi-instance workloads.

## 4 Conclusion and Discussion

In this work, we demonstrated that modern CPU architectures, when coupled with hardware-aware algorithmic optimizations, are not only viable but superior for end-to-end protein structure prediction pipelines. By addressing the architectural mismatch between AlphaFold2’s preprocessing and attention modules and traditional GPU acceleration, we achieved a transformative reduction in execution time. Our “key” results — (1) for a 2752-length protein, reducing the preprocessing latency from approximately 90 mins to just under 12 mins and the inference latency from nearly 69 hours to just 2.7 hours on CPU and (2) demonstrating 76% higher end-to-end throughput for a 391-protein proteome using a dual-socket Intel Xeon 6980P system compared to a configuration with single CPU socket with H100 GPU offload — highlight the potential of Intel Xeon 6980P for large-scale biological research. The shift from days to hours for complex structures, combined with nearly 97% and 83% reduction in memory footprint for gating attention and triangle multiplication, respectively, provides a scalable and cost-effective alternative to GPU-restricted workflows, paving the way for proteome-wide structural analysis on versatile, ubiquitous CPU-based infrastructure.

In this work, we deliberately chose AlphaFold2 for acceleration instead of the newer AlphaFold3. While AlphaFold3 inference code has been made publicly available, its restrictive commercial licensing has limited its adoption compared to the ubiquity of AlphaFold2. Given AlphaFold2’s significantly larger user base and its status as the current industry standard for open-source workflows, it remains the primary target for hardware optimization. Furthermore, because AlphaFold2 and AlphaFold3 share several core computational modules, the acceleration strategies developed in this work — specifically our efficient Gating Attention and Triangle Multiplication kernels — are directly transferable to the AlphaFold3 architecture. For instance, the Pairformer stack in AlphaFold3 employs similar attention-driven geometric refinement logic, and even the newer Diffusion modules can leverage our tiling and fusion principles. Thus, the architectural principles established here provide a clear, high-performance roadmap for a future acceleration of AlphaFold3.

The performance improvement shown in this paper stems from several factors that arise in end-to-end, real-world applications:

(1) In many such applications, deep learning constitutes only a portion of the overall computation. Other components – such as database searches, string matching, and graph operations – account for a significant fraction of the application execution time. These tasks are often irregular and difficult to vectorize efficiently on GPUs, limiting their acceleration potential.
(2) Even within the deep learning component, many operations exhibit low arithmetic intensity. For instance, in AlphaFold2, the dimensions of matrices for matrix multiplication are determined by protein sequence length, number of attention heads and head sizes. Given the smaller number of attention heads and smaller head sizes in AlphaFold2, the matrix multiplications lack sufficient arithmetic intensity to fully utilize the parallel processing capabilities of modern GPUs. The situation is even worse for shorter proteins.
(3) GPU kernel launch overhead can also be a significant cost. While this is negligible for large, long-running kernels, it can become a bottleneck for smaller kernels, reducing effective throughput.

CPU platforms have historically excelled at handling irregular computational workloads. Newer CPU generations also featuring enhanced matrix multiplication capabilities – enabled by AMX type instructions. A well-optimized, scalable software that leverages both the flexibility for irregular compute and the accelerated matrix multiplication capabilities of modern CPUs can serve as a compelling alternative to GPU-based computing for end-to-end inference tasks. In many scenarios, a CPU with robust deep learning support may emerge as a faster, more efficient, and cost-effective solution making it a strong contender for the future of AI inference.

## 5 Methods

In Section 3.3, we provided a brief overview of our acceleration techniques including multi-processing with dynamic scheduling, acceleration of preprocessing through enhanced vectorization and use of optimal compilation and execution setup, acceleration of model inference through cache optimizations, operator fusion and use of reduced precision (BFloat16) that also enabled the use of AMX. In this Section, we provide a detailed description of our acceleration techniques for model inference. In Section 3.2, we demonstrated that the majority of inference time is spent in two generalized modules – *Gating Attention* and *Triangle Multiplication*. In this Section, we describe our acceleration of these two modules.

### Algorithm 1

Gating Attention Algorithm. Illustrated without batch dimension (*B*) looping for simplicity.

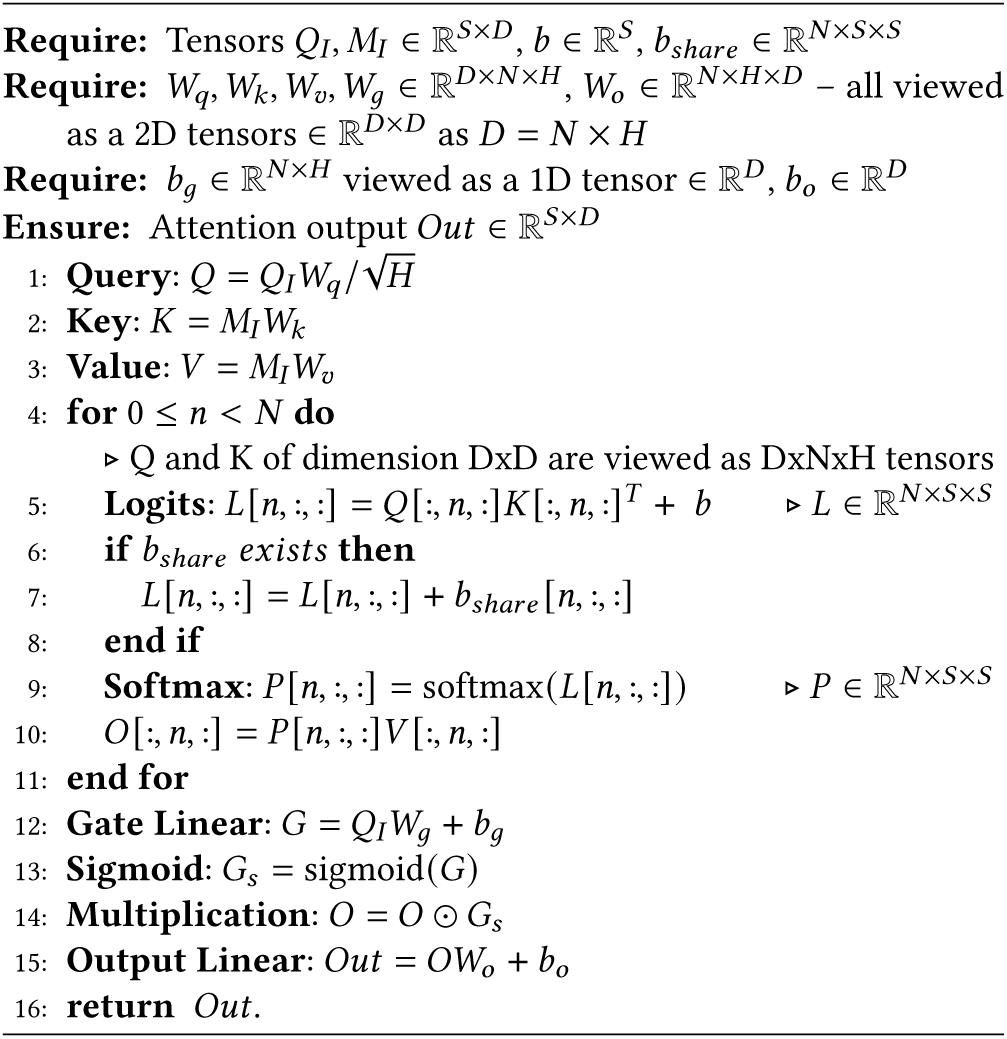

### 5.1 Notation

For two matrices *X* and *Y*, we use *XY* to denote their multiplication, *X + Y*to denote their element-wise addition and *X Y* to denote their element-wise multiplication. For a matrix *X* and a vector *b*, *X + b* denotes addition of each element of *b* to all the elements of the corresponding column of *X* and *X b* denotes multiplication of each element of *b* to all the elements of the corresponding column of *X*. We employ Python-style indexing for tensors, so *X* 3, : refers to the third row of the two dimensional tensor *X*. Unless stated otherwise, all tensor operations are implicitly batched. Matrix products such as *XY* represent batched matrix multiplication over the leading batch dimension, which is omitted from the notation for brevity.

### 5.2 Baseline Gating AEention and Triangle Multiplication

#### 5.2.1 Baseline Gating AFention

Gating attention is a generic multi-headed cross-attention module. It is invoked within several modules of AlphaFold2 and AlphaFold2-Multimer with varying parameters and input tensor dimensions. Algorithm 1 illustrates the gating attention module. The module modifies the input embedding by first applying multiple projections (query, key, value) and then computing scaled dot-product attention (SDPA) using multiple attention heads. It takes four input tensors: a queries tensor (*Q_I_*), a memories tensor (*M_I_*), a bias tensor (*b*), and a shared bias tensor (*b_share_*). The queries (*Q_I_*) and memories (*M_I_*) tensors have dimension *B S D*, where, *B* is the batch dimension, *S* is the sequence dimension along which attention is applied, and *D*is the embedding dimension. The bias and shared bias tensors have dimensions *B × S* and *N S S*, respectively, where *N* is the number of attention heads. The shared bias is shared across all batches. In addition to the input tensors, the gating attention layer contains several weights parameters (*W_q_, W_k_, W_v_, W_g_, W_o_*) for the linear layers. The input layer weights (*W_q_, W_k_, W_v_, W_g_*) have dimensions *D NH*, where *H* is the head dimension. The output layer weight (*W_o_*) has dimensions *N H D*. Both the gate and output linear layer weight parameters also include a bias term (*b_g_, b_o_*).

To give an intuition of how Gating Attention module is used to implement different attention modules, here are a few examples. Gating attention module implements cross attention, and therefore, uses separate *Q_I_* and *M_I_* tensors. To use it for self attention, we set *M_I_* = *Q_I_*. In MSA row-wise self attention module, attention is applied to each MSA row independently. In this case, Gating attention is used with batch size (*B*) equal to the number of MSA rows and sequence length (*S*) equal to the length of the protein sequence. On the other hand, in column-wise self attention, where attention is applied to each MSA column independently, *S*is the number of MSA rows and while *B* is the protein sequence length. In case of pair representation, the number of rows and columns are both equal to the protein sequence length. Therefore, *B*= *S* = protein sequence length.

#### 5.2.2 Baseline Triangle Multiplication

Triangle multiplication module updates the pair representation in the Evoformer block, providing a mechanism for the network to enforce the triangle inequality in the triangles of the 3D structures. For each triangle, it combines information from two of its edges to update the third edge. The module has two separate forms, one for “outgoing” edges and one for “incoming” edges. However, both forms can be captured in a single algorithm by using different equations for “incoming” and “outgoing” edges. The original AlphaFold2 code used a version of triangle multiplication module that processed the left edge and right edge separately. However, in AlphaFold2-Multimer, a newer fused triangle multiplication version has been used that processes both of them together. We have optimized both versions using the same techniques, but here we only discuss the fused version (Algorithm 2) as it is easier to understand.

The fused triangle multiplication module takes the pair representation *Z* as input and produces an updated version of it through a series of transformations. The pair representation tensor *Z* has dimensions *S S D*, where S is the sequence length, equal to the length of the protein, and *D*is the embedding dimension. This pair representation tensor undergoes multiple linear projections and gating operations, which transform the hidden dimension of intermediate tensors from *D* to *C*. Subsequently, the left and right projection tensors are multiplied according to either the “incoming” or “outgoing” edge formulation. Finally, additional linear projections and gating operations are applied to transform the hidden dimension back from *C*to *D*. The module also includes auxiliary operations such as layer normalization, sigmoid activation, and element-wise multiplication applied to the intermediate tensors.

##### Algorithm 2

Fused Triangle Multiplication

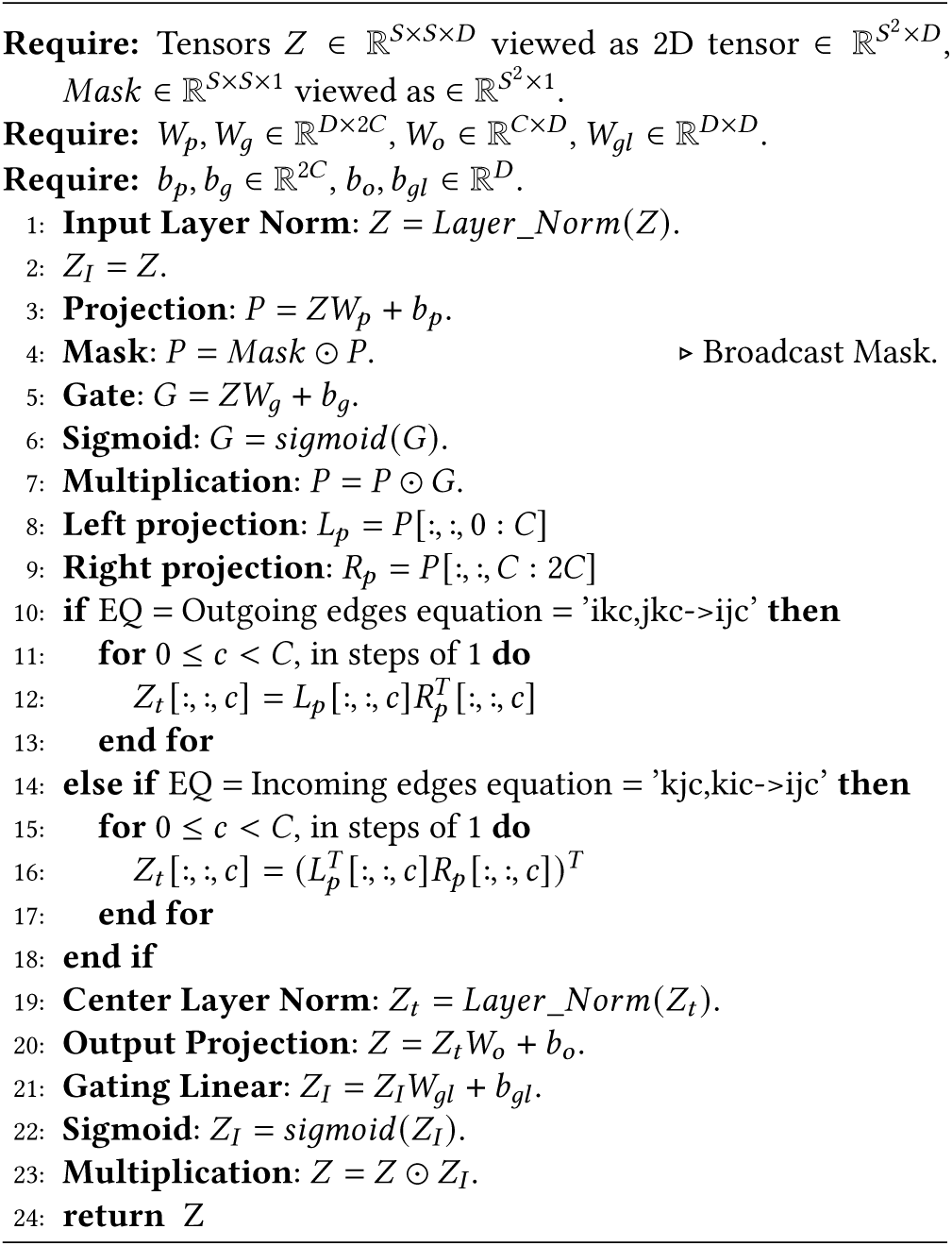

#### 5.2.3 Memory Capacity, Bandwidth and Compute Requirements

In each computation step of Gating attention (Algorithm 1), reading and writing of one or more large tensors from memory is needed. In particular, intermediate tensor *L*and *P* have large dimensions of *B N S S*. In the row-wise attention module, the number of attention heads (*N*) is 8, the batch size (*B*) corresponds to number of rows in MSA representation and the sequence length (*S*) corresponds to the input protein length. This means that memory capacity requirements and inference time increases quadratically with the input protein length. Lengths of protein found in nature range in a few hundreds to a few thousands with a majority of the proteins having length less than 1000. The number of MSA rows depends on number of matches found and is capped at 512 in AlphaFold2. With these values, the memory requirements for just tensor *L* can be several GBs. The combined memory requirement of all the tensors can easily range in several tens of GBs and may even be a few hundred GBs for longer proteins.

These tensors do not fit in caches, a problem that becomes more pronounced as the sequence length (*S*) increases. Therefore, in a naive implementation, they need to be read from memory for every operation (e.g. computation of Q, K and V, softmax, sigmoid, etc.), resulting in the number of bytes read from the memory for each operation of the order of the size of the tensors. This makes these operations memory bandwidth-bound. For matrix multiplication based operations, given that the number of heads (*N* = 4, 8) and head sizes (*H* = 8, 16, 32) are comparatively smaller than in the multi head attention (MHA) of standard LLMs, it reduces their arithmetic intensity (the ratio of computational FLOPS required to bandwidth required – also known as flops-per-byte ratio). This is due the fact that embedding dimension *D* of transformer is *N ×H*, and flops-per-byte ratio is directly proportional to the embedding dimension. Additionally, the element-wise operations, such as, softmax, sigmoid, addition, element-wise multiplication etc., have computational FLOPS required also of the order of the size of the tensors. Therefore, the flops-per-byte ratio of these operations is even lower than that of matrix multiplication. In practice, the flops-per-byte ratio of all of the above operations (matrix multiplication and element-wise) is significantly smaller than that of the modern CPUs/GPUs (See Tables S3 and S4), making these operations primarily bandwidth-bound.

Moreover, with a naive implementation, the memory capacity requirement may be higher than available memory on GPU or even a CPU (as exemplified by OOM errors for baseline implementation in Table 6). Therefore, techniques that can reduce the memory capacity and bandwidth requirement should help in accelerating this module.

Similarly, for Fused triangle multiplication (Algorithm 2), each step requires reading and writing large tensors from memory. The sizes of these tensors increase quadratically with the sequence length *S*. Many steps in the algorithm are also element-wise operations, similar to those in the gating attention module. Consequently, the triangle multiplication module is also bandwidth-bound.

### 5.3 Prior Acceleration Efforts

For GPUs, several acceleration techniques [19–22] have been proposed to alleviate the bandwidth and memory capacity constraints of the standard multi-head attention (MHA) module. Projection layers are standard linear layers without bias and, therefore, can leverage preexisting optimized GPU implementations for the linear layer. In accelerated GPU implementations of SDPA, computation is parallelized across the number of heads and along the batch dimension. This approach works reasonably well with smaller sequence lengths due to the high bandwidth memory (HBM) of GPUs. However, at longer sequence lengths, that increases the bandwidth pressure, even HBM equipped GPUs face memory bandwidth constraints. Moreover, GPUs also face capacity constraints due to large tensor sizes and limited sizes of HBM. Therefore, it becomes imperative to utilize memory tiling along the sequence dimension and operator fusion as used by FlashAttention [20], FlashAttention-2 [21], and FlashAttention-3 [22].

#### Tiling, Online Softmax, Operator Fusion and Parallelization

In tiling, input sequences are partitioned into smaller contiguous blocks (called tiles) such that the tiles and corresponding intermediate tensors of these tiles fit into caches. Each tile is processed independently while performing multiple operations (e.g. computing *QK^T^*, softmax, and *PV*) reusing the tiles and intermediate tensors from cache multiple times before moving to the next tile. The approach of reusing a tile that is loaded in cache multiple times is called tiling, while the approach of performing multiple operations on each tile, rather than writing the tensors to memory after each operation and reading the same tensors from memory before the next operation, is called operator fusion. Tiling needs adjustments in the softmax algorithm. Softmax is performed on a row of a tensor and requires the row’s maximum value for computation. When processing a tile, the entire row is not available to get the maximum value. Online softmax technique [23] allows partial softmax computation on a tile via use of local maximum and fixing up of values later as needed. These approaches reduce repeated reads of large sequence tensors from HBM, improve cache locality, and lower memory bandwidth pressure, which is especially important for long protein sequences. These techniques are briefly illustrated in Figures 4(c) and 4(d) where one-dimensional and two-dimensional tiling (with partial softmax) is performed and "bias" and "softmax" operations are fused. Tiling creates more opportunities for parallelism; you can process individual tiles simultaneously, alongside the existing parallelism in the batch and head dimensions. In Supplementary Section S3.3, we provide a more detailed explanation of memory tiling, online softmax, operator fusion and multi-threading based parallelization.

In comparison to GPUs, there have only been a few efforts to accelerate the MHA module for CPUs. CPUs face even higher band-width constraints because they employ DRAM that has lower band-width compared to HBM. A previous attempt to accelerate the MHA module on CPUs [24] utilized five-dimensional tiled layout tensors throughout the network. Here, layout of embedding tensor was converted from three dimensions *B S D* into five dimensions by tiling along the S and D dimensions. This work employed tiling for acceleration but it didn’t use the online softmax [23] approach used in FlashAttention-2,3 algorithms. Additionally, this approach used the same tiled tensor representation throughout the network, which is less suitable for the gating attention module of AlphaFold2 and AlphaFold2-Multimer. Firstly, in AlphaFold2 model inference, the gating attention sequence length (*S*), number of heads (*N* = 4, 8) and head size (*H* = 8, 16, 32) are not fixed for the entire network. Secondly, row-wise and column-wise attention have different sequence lengths. Sequence length also depends on the input protein length for some sub-modules. Consequently, input tensors cannot be tiled assuming a fixed sequence length, number of heads, or head size. Additionally, the small number of heads and small head sizes prevent tiling along these dimensions while still maintaining computational efficiency. Therefore, we implement a multi-headed attention module, where we tile only along the sequence length dimension. This allows our implementation to use the same module to handle any combination of sequence length, number of heads and head size appearing in AlphaFold2 and AlphaFold2-multimer.

### 5.4 Our Acceleration Approach

Prior approaches, such as FlashAttention and FlashAttention-2 [20, 21], parallelize computation across the batch and head dimensions. However, this strategy is inefficient for AlphaFold2’s gating attention module, where the head count is low (*N* = 4 or 8). This inefficiency is further exacerbated by the increasing core counts of modern CPUs, which leads to significant hardware underutilization. For accelerating the Gating attention and Triangle multiplication modules, we primarily use tiling along the sequence length dimension and operator fusion to improve cache efficiency. This also creates more opportunities of parallelism – you can process individual tiles simultaneously – that we exploit by multi-threading across tiles in addition to multi-threading across the batch and head dimensions. Table 9 provides a brief summary of acceleration techniques used for various kernels. We use modules from LIBXSMM library to perform tile-level operations on CPU at high efficiency. As we discuss in Supplementary Section S3.4, LIBXSMM provides highly optimized, low-level deep learning primitives – both individual and fused – for CPU architectures. By abstracting hardware-specific implementations, such as Intel AMX instructions, it offers a high-level tensor API, supporting operations such as matrix algebra and element-wise kernels.

**Table 9:** Optimization Techniques Used in Various Kernels.

| Module | Kernel | Techniques |  |  |
| --- | --- | --- | --- | --- |
|  |  | Tiling | Multi-threading | Operator Fusion |
| Gating Attention | Projection | 1D | Yes | Yes |
|  | SDPA | 2D | Yes | Yes |
|  | Output | 1D | Yes | Yes |
| Fused Triangle Multiplication | Proj | 2D | Yes | Yes |
|  | Eqn | 2D | Yes | Yes |
|  | Out GEMM | 1D | Yes | Yes |
|  | Gate GEMM | 1D | Yes | Yes |

### 5.5 Our Acceleration of Gating Attention

We divide the gating attention module into three steps - projection (lines #1-3 and transpose of *K*), SDPA (lines #4-11) and output (lines #12-15). In the following, we provide details on acceleration of each of these steps.

#### 5.5.1 Projection Step

We write three similar kernels for creating projections of *Q*, *K*and *V* tensors. All three kernels, multiply the corresponding tensors and weight matrices - essentially performing matrix multiplication. We fuse the matrix multiplication with multiplication by 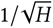 for *Q* and transpose for *K*. In these kernels, we tile both the input and output tensors along their sequence length dimension (*S*). We parallelize the computation across tiles and along the batch dimension using multi-threading. For each tile of the input tensor, we multiply it with the weight matrix, perform additional operations (multiplication, transpose) if any, and eventually write to a tile of the output tensor. Algorithm S1 illustrates this with the kernel for *K* tensor.

#### 5.5.2 SDPA Step

We fuse the lines #4-11 of Algorithm 1 and refer the fused kernel as *Scaled dot product attention (SDPA)*. Our *SDPA* is similar to the SDPA operation inside the MHA module of modern LLMs with some additional bias additions.

##### Operator Fusion & Tiling

We implement two separate tiling approaches for the *SDPA* kernel and choose the suitable approach based on the input protein sequence length and compute precision. In the first algorithm (Algorithm S2, Figure 4(c)), we tile only Q along the sequence dimension and perform full softmax computation. This algorithm is suitable for short sequences, especially when all the intermediate data corresponding to a tile of *Q* can be kept in CPU caches. Second algorithm (Algorithm S3, Figure 4(d)) uses FlashAttention-2 approach where we tile all of *Q* and *K^T^* and *V*along the sequence dimension and perform partial softmax computation on tiles. This approach reduces the size of the intermediate data at the cost of additional logic and data for partial softmax and later fixup and is more suitable for longer sequences for which full softmax computation becomes inefficient due to cache misses. In Algorithm S2, for each tile of Q, we perform matrix multiplication with *K^T^* and use an intermediate buffer of size *T_q_ S*to store the output, where *T_q_* is the tile size. Subsequently, we add tiled biases and perform full softmax computation on this buffer followed by multiplication with *V*. This algorithm has an additional advantage that it enables several consecutive matrix multiplications with the same dimensions, thereby, minimizing the number of times the matrix multiplication hardware must reconfigure its settings, leading to much higher efficiency. In Algorithm S3, we tile all of *Q*, *K^T^* and *V* along the sequence dimension leading to a smaller intermediate buffer of size *T_q_ T_kv_*, where *T_q_* is the tile size for *Q* and *T_kv_* is the tile size for *K^T^* and *V*. We perform matrix multiplication on tiles of *Q* and *K^T^*, add tiled biases, do partial softmax computation, and eventually multiply by the corresponding tiles of *V*. In the end, we perform a fixup of the softmax values. We choose the value of *T_kv_*based on a tradeoff. The value needs to be small enough for intermediate buffer to fit in cache. However, anytime we switch between multiplications of matrices of different dimensions (line #7 and #13), the matrix multiplication hardware must reconfigure its settings. Therefore, to minimize the reconfigurations, we need to use as large a *T_kv_* as possible.

Therefore, our *SDPA* kernels fuse multiple operations, performing them on tiles that are maintained in cache and write tiles to memory only after all the computations are performed. This reduces round trips to memory, thereby alleviating bandwidth bottlenecks. We also reduce the need for storing large tensors, thereby reducing memory capacity requirements for inference on long protein sequences.

##### Multi-threading

We parallelize the computation across tiles and across batch and head dimensions using multi-threading.

##### Overlapping matrix multiplication with element-wise operations

On modern processors, there are dedicated matrix engines – such as GPU Tensor cores or CPU AMX units – that perform matrix multiplication, while the element-wise operations – e.g. softmax – is performed on SIMD/SIMT units. To achieve high efficiency, the dedicated matrix engines must remain active as often as possible. However, the FlashAttention-2 algorithm can become inefficient when the ratio of matrix multiplication to softmax compute is low, as the matrix engine sits idle while the SIMT units process the soft-max. FlashAttention-3 [22] mitigates this on GPUs by overlapping matrix multiplication and partial softmax via complex asynchronous instructions. CPUs with AMX units face a similar bottleneck; however, we resolve this by leveraging hyper-threading. By running two threads per core – one dedicated to matrix multiplication and the other to softmax – the CPU hardware naturally overlaps these tasks. This provides a “FlashAttention-3 style” capability without requiring complex algorithmic redesign.

The effectiveness of this overlap depends on the attention head size. In the Attention module, a larger head size increases the ratio of matrix multiplication to softmax computation. While most Large Language Models use high head sizes (*H* ≥ 64) to balance these operations, AlphaFold2 utilizes much smaller head sizes (8, 16, 32). This creates a significant compute imbalance that prevents GPUs from fully utilizing their high matrix engine FLOPS, showing that GPUs high FLOPS are surplus to the requirements of AlphaFold2. Consequently, the Xeon CPUs with lower peak throughput of the matrix engine still perform competitively with the GPUs.

#### 5.5.3 Output Step

In the final kernel, we fuse the lines #12-15 of Algorithm 1 and call the fused kernel as the *output* kernel (Algorithm S4). In this kernel, we again tile along the sequence dimension. We fuse two linear layers and gate multiplication operation in this kernel and only store the final results to memory thereby reducing memory bandwidth requirements.

### 5.6 Acceleration of Triangle Multiplication

Similar to the gating attention module, we use a tiling based approach to accelerate the triangle multiplication module. However, unlike the gating attention module, we not only tile along both sequence dimensions but also perform data layout transformation using permute operation for better cache locality. First, we divide the sequence dimension (*S*) into 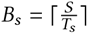 tiles of size *T*_s_. Later, we convert three-dimensional intermediate*^Ts^*tensors of dimension *S S C* into a five dimensional tensors of dimension *C B_s_ B_s_ T_s_ T_s_*. This tiling strategy enables better cache locality and allows us to use the LIBXSMM primitives more efficiently.

We divide the triangle multiplication module (Algorithm 2) into three steps - projection (lines #3-9), equation (lines #10-18) and output (lines #19-23). In the following, we provide details on acceleration of each of these steps.

#### 5.6.1 Projection Step

We fuse lines #3-9 of Algorithm 2 and refer to that as *projection* kernel. In this kernel, we tile along the sequence dimension and fuse multiple linear layers, gating and transpose operations to create left projection and right projection tensors (Algorithm S5). For each tile, we read it from memory and the final results are stored to memory only when all the compute steps have finished. Additionally, transposed weight and bias tensors 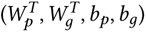 are kept in cache due to their small size. We parallelize the computation across tiles using multi-threading.

#### 5.6.2 Equation Step

We combine steps #10-18 of Algorithm 2 and refer to the fused kernel as the equation kernel (Algorithm S6). This kernel takes the projection outputs from the *projection* kernel (*L_p_, R_p_*) and implements EINSUM equations specific to incoming edges and outgoing edges. We tile along the sequence dimensions and use LIBXSMM primitives which enable efficient matrix multiplication of multiple blocks. At the end of this computation, we permute five dimensional tensors and reshape them to convert back into two three dimensional tensors. We parallelize the computation across tiles and along the embedding dimension using multi-threading.

#### 5.6.3 Output Step

In addition to the previous kernels, we also fuse steps #19-23 and call it the *output* kernel. The structure of this *output* kernel is similar to the *output* kernel of gating attention module in that it combines linear layers with operations such as *multiplication* and *sigmoid*. Therefore, we use the same template as was used for linear layers in Algorithm S4 and, for the sake of brevity, skip discussion of this kernel.

### 5.7 Implementation with BF16 Precision

While the original AlphaFold2 implementation relies on single-precision (FP32), modern deep learning frameworks like PyTorch support reduced-precision tensors and arithmetic. In bandwidth-bound workloads, BFloat16 can provide up to a 2x speedup, while the performance gains in compute-bound scenarios can be even more significant. Consequently, we implemented Evoformer inference in BFloat16 to leverage the full architectural capabilities of modern CPUs. This transition involved casting input tensors to BFloat16 and ensuring that all successive computational steps maintain this precision.

However, reducing precision within PyTorch does not automatically guarantee efficient execution. Transitioning to BFloat16 is just a prerequisite for utilizing Intel AMX on Xeon CPUs. To enable efficient execution, we optimized the gating attention and triangle multiplication layers to specifically target AMX instructions. We leveraged primitives from the LIBXSMM library, which interface seamlessly with reduced-precision data. By employing BFloat16-specific primitives, we adapted the optimized algorithms described in the previous section to generate Intel AMX instructions that utilize the hardware’s matrix engines. While converting these kernels from FP32 to BFloat16 requires minor structural adjustments—such as VNNI (Vector Neural Network Instructions) reformatting and precision casting for element-wise operations—we omit these implementation details for brevity.

## Acknowledgments

We extend our gratitude to the DeepMind AlphaFold team for their groundbreaking contribution to the field and for making their model parameters and source code accessible to the scientific community. We also thank the developers of the LIBXSMM library, particularly for their assistance in implementing the Tensor Processing Primitives (TPPs) that serve as the foundation for our optimized Gating Attention and Triangle Multiplication kernels. Finally, we acknowledge the contributors to the Open-Omics Acceleration Framework, whose collaborative efforts continue to drive the optimization of life sciences pipelines.

We used large language models to help polish the writing. We take full responsibility for all content in this paper.

Optimization Notice: Software and workloads used in performance tests may have been optimized for performance only on Intel microprocessors. Performance tests, such as SYSmark and MobileMark, are measured using specific computer systems, components, software, operations and functions. Any change to any of those factors may cause the results to vary. You should consult other information and performance tests to assist you in fully evaluating your contemplated purchases, including the performance of that product when combined with other products. For more information go to http://www.intel.com/performance. Intel, Xeon, and Intel Xeon Phi are trademarks of Intel Corporation in the U.S. and/or other countries.

## S1 Supplementary Tables

**Table S1:**
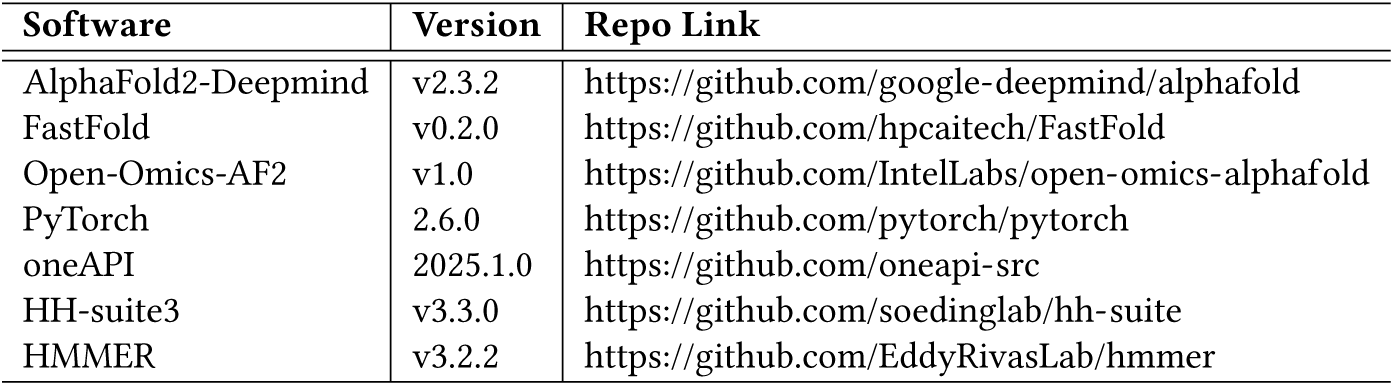
Software and Codebase versions.

| Software | Version | Repo Link |
| --- | --- | --- |
| AlphaFold2-Deepmind | v2.3.2 | https://github.com/google-deepmind/alphafold |
| FastFold | v0.2.0 | https://github.com/hpcaitech/FastFold |
| Open-Omics-AF2 | v1.0 | https://github.com/IntelLabs/open-omics-alphafold |
| PyTorch | 2.6.0 | https://github.com/pytorch/pytorch |
| oneAPI | 2025.1.0 | https://github.com/oneapi-src |
| HH-suite3 | v3.3.0 | https://github.com/soedinglab/hh-suite |
| HMMER | v3.2.2 | https://github.com/EddyRivasLab/hmmer |

**Table S2:** Genetic (sequence) databases.

| Dataset | Repo Link |
| --- | --- |
| BFD | https://bfd.mmseqs.com/ |
| MGnify | https://www.ebi.ac.uk/metagenomics/ |
| PDB70 | http://wwwuser.gwdg.de/~compbiol/data/hhsuite/databases/hhsuite_dbs/ |
| PDB | https://www.rcsb.org/ |
| PDB seqres | https://www.rcsb.org/ |
| UniRef30 | https://uniclust.mmseqs.com/ |
| UniProt | https://www.uniprot.org/uniprot/ |
| UniRef90 | https://www.uniprot.org/help/uniref |
| Proteomes Caenorhabditis elegans | https://www.uniprot.org/proteomes/UP000001940 |

**Table S3:**
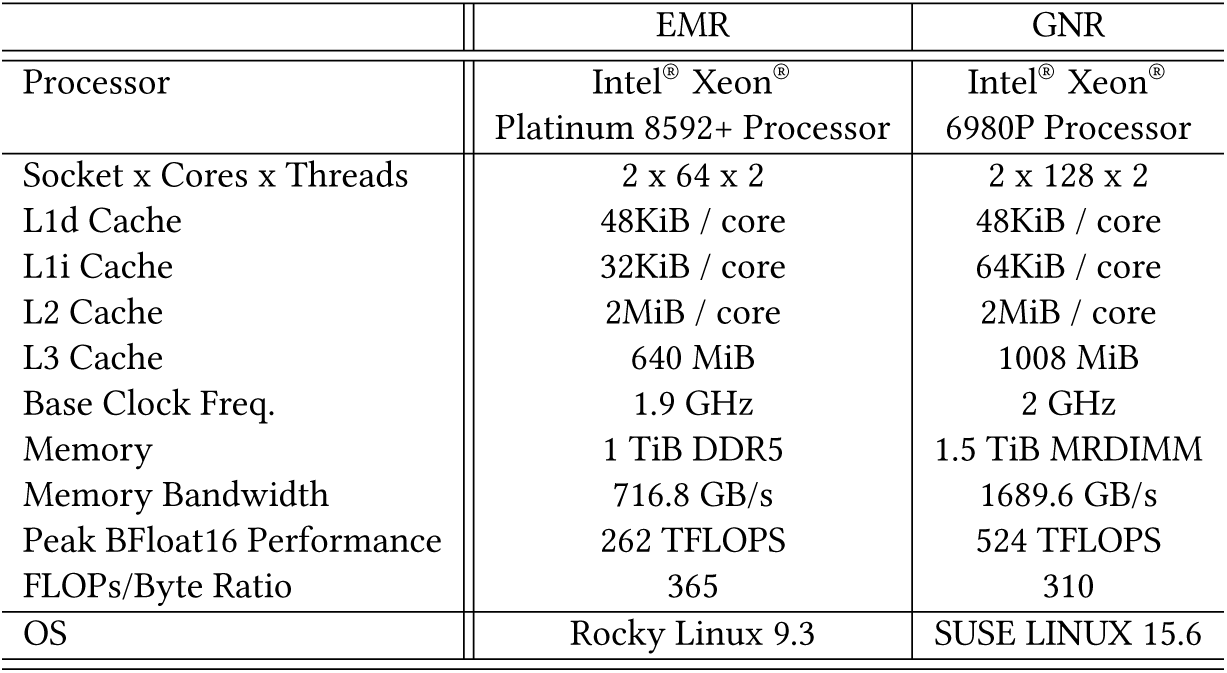
Details of the CPU systems used in our experiments.

**Table S4:** Details of the GPU systems used in our experiments.

|  | A100 | H100 |
| --- | --- | --- |
| GPU | Nvidia A100 40 GB | Nvidia H100 80 GB |
| Instance | GCP a2-highgpu-1g | On-Prem |
| Memory | 40 GB HBM2 | 80 GB HBM3 |
| Memory Bandwidth | 1555 GB/s | 3350 GB/s |
| Peak TF32 Performance | 156 TFLOPS | 495 TFLOPS |
| Peak BFloat16 Performance | 312 TFLOPS | 990 TFLOPS |
| FLOPs/Byte Ratio | 200 | 295 |
| OS | Ubuntu Linux 22.04 | SUSE Linux 15.6 |

**Table S5:** Tensor processing primitives (TPP) [24] used in this paper.

| TPP | Operation | Inputs | Outputs |
| --- | --- | --- | --- |
| GEMM | Matrix Multiplication | $A, B$ | $A * B$ |
| BIAS | Bias Addition | $A, b$ | $A + \text{Broadcast}(b)$ |
| ADD | Addition | $A, B$ | $A + B$ |
| MUL | Multiplication | $A, B$ | $A \odot B$ |
| TRANS | Transpose | $A$ | $A^T$ |
| SOFTMAX | Softmax | $A$ | $\text{Softmax}(A)$ |
| SOFTMAX_PARTIAL | Partial Softmax on a tile | $A$ | $\text{Softmax}(A), \text{max}, \text{sum}$ |
| SOFTMAX_FIXUP | Partial Softmax fix<br>from current maximum value | $A, C_{\text{max}}, C_{\text{sum}}, O_{\text{max}}, O_{\text{sum}}$ | $A, O_{\text{max}}, O_{\text{sum}}$ |
| SOFTMAX_FLASH_SCALE | Final softmax scaling | $A, O_{\text{sum}}$ | $A$ |
| SIGMOID | sigmoid | $A$ | $\text{Sigmoid}(A)$ |
| MUL_BCAST | Broadcast Multiply | $A, b$ | $A \odot \text{Broadcast}(b)$ |
| BRGEMM | Batch reduce GEMM | $A, B, A_{\text{stride}}, B_{\text{stride}}$ | $C = \sum_i A_i * B_i$ |

## S2 Supplementary Figures

**Figure S1:**
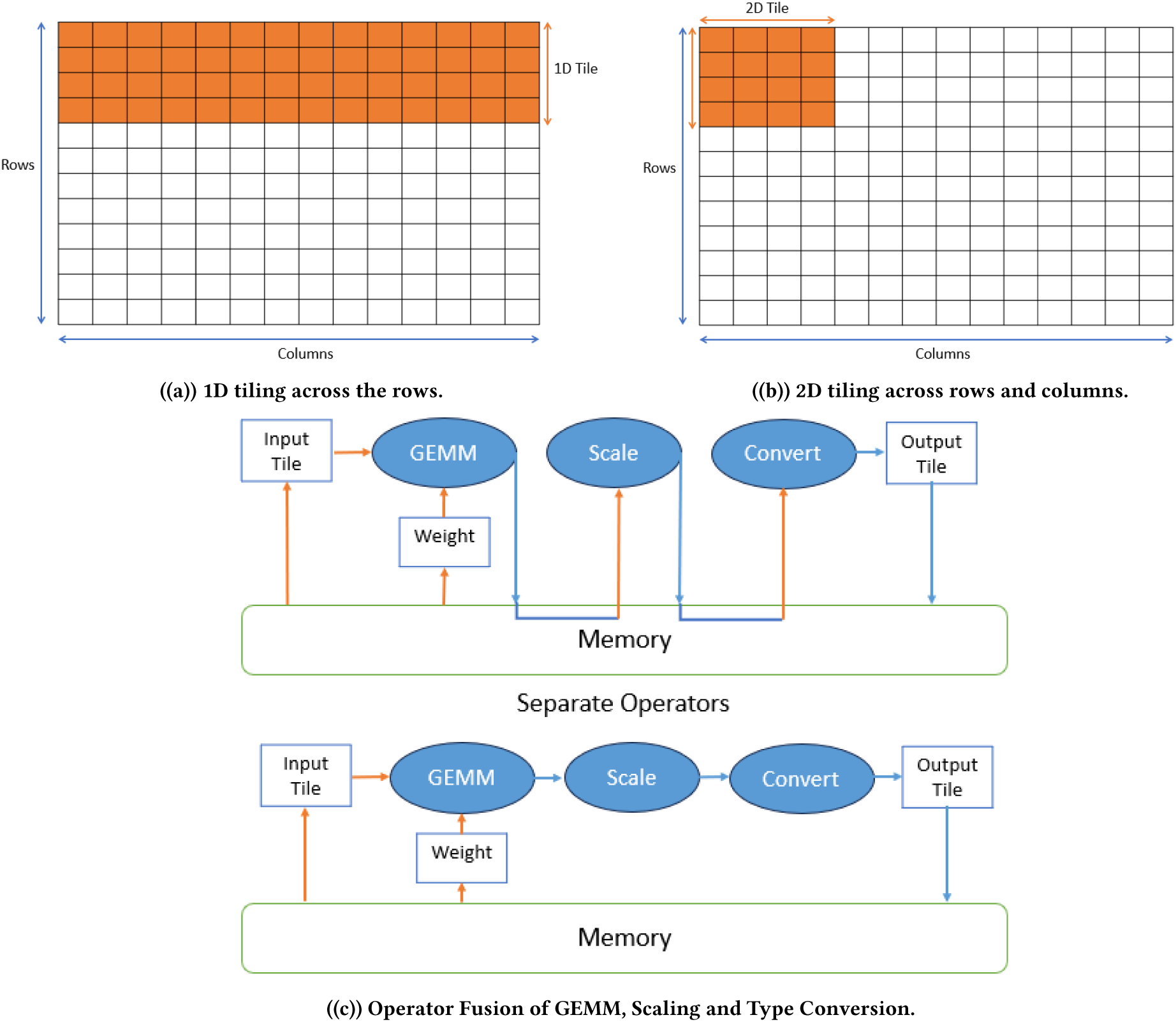
Illustration of Key Kernel Optimization techniques.

## S3 Supplementary Text

### S3.1 Multi-Processing with Dynamic Scheduling

We divide the folding of the input proteins into two phases. In the first phase, we perform preprocessing of all the input proteins and store the intermediate results to disk. In the second phase, we use the intermediate results from preprocessing to perform model inference on all the proteins.

For each of the above phases, we use multi-processing with dynamic scheduling. In the following description, we define a task as preprocessing or inference of a single protein. Given a set of tasks, we process them as follows. We sort the tasks in decreasing order of expected run time. For example, for inference phase, we sort the inference tasks in decreasing order of protein lengths. Based on the memory capacity, expected memory required for the tasks and number of tasks, we calculate the number of parallel tasks, say *n*, and number of cores per task. We begin by launching the first *n* tasks from the sorted list in parallel assigning equal number of cores to each of them. When a task is completed, the corresponding cores are immediately assigned the next one from the sorted list to ensure that cores do not sit idle. If a task fails due to out-of-memory (OOM) error, it is added to a list of such tasks. We attempt to process them in the next round by reducing the number of parallel tasks, thereby, increasing the memory available per task. This is repeated, reducing the number of parallel tasks in each round, until all the tasks are processed. Sorting of tasks in descending order of expected execution time ensures that the longest tasks get more tasks to overlap with. This dynamic scheduling approach allows us to minimize idling of CPU cores.

### S3.2 Optimizations to accelerate preprocessing

We performed a series of software optimizations on the preprocessing tools. Our optimization strategy for preprocessing primarily involved configuration adjustments, compiler optimizations, AVX-512 based vectorization, and parallel execution tuning.

#### S3.2.1 Optimization of HHblits

We accelerate HHblits through the optimization of the execution setup. The HHblits uses OpenMP-based multithreading and is characterized by several small memory allocations. We fine-tune OpenMP performance by using the following KMP_AFFINITY setting:

`KMP_AFFINITY=granularity=fine,compact,1,0`

This substantially increased the efficiency of OpenMP multi-threaded execution. To optimize memory management, we integrate the jemalloc library, resulting in a significant performance enhancement. The combination of these optimizations have significantly elevated the overall performance of HHblits.

#### S3.2.2 Optimization of Jackhmmer

Through VTune profiling, we identified the hotspot of Jackhmmer as "calc_band_12" that accounted for a significant portion of execution time. The original implementation of this function relied on 256-bit AVX based vector instructions. In order to optimize its performance, we re-implemented it using 512-bit AVX512 instructions. To further augment the performance, we replaced GCC compiler with ICC (Intel C/C++ Compiler). These optimizations significantly contributed to enhancing the overall performance of Jackhmmer.

#### S3.2.3 Optimization of HHSearch

The original HHSearch uses 128-bit SSE4 based instructions for vectorization. We re-implemented it with hand-written 512-bit AVX512 based vector instructions. This achieved nearly a 33% performance boost. We further improved the performance through compiler optimizations. We applied a combination of compiler flags, including `-O2/-O3, -march=sapphirerapids, -flto, -finline-functions`, and `-funroll-loops` to provide the compiler with additional hints to generate optimized binary code. These modifications resulted in an approximate 25% speedup.

### S3.3 Explanation of Acceleration Techniques Used for Inference

#### S3.3.1 Tiling

In modern CPU systems, memory system is significantly slower compared to the CPU’s arithmetic units. Therefore, CPUs employ multiple levels of high bandwidth cache to mitigate this performance bottleneck and keep the CPU busy during computations. However, these caches are typically small in size, which means that operations involving large matrices can easily exceed cache capacity, leading to frequent and costly memory accesses. Tiling is a widely used optimization technique to address this issue. It involves breaking down matrix operations into smaller tiles that are small enough to fit within the cache, so that data can be reused efficiently while minimizing memory traffic. Tiling can be applied in one dimension (1D tiling) or both dimensions (2D tiling). For instance, 1D tiling might involve processing 16 rows of a matrix across all columns at once, while 2D tiling could process a block of 16 rows by 64 columns simultaneously. Figure S1 illustrates examples of both 1D and 2D tiling strategies. In the context of AlphaFold2, the model’s embedding dimensions and weight matrix dimensions are relatively small (e.g., 64, 128, 256). Given these constraints, we apply tiling only along the protein sequence length dimension in our kernels for AlphaFold2.

#### S3.3.2 Operator Fusion

is another technique technique designed to make more efficient use of the cache hierarchy by minimizing expensive memory accesses. Since reading from and writing to memory can be costly in terms of performance, operator fusion aims to load a data tile from memory only once, apply multiple operations to it while it resides in faster cache or registers, and then write it back once the computations are complete. This approach significantly reduces memory bandwidth pressure. Figure S1 gives an example of operator fusion that combines GEMM with scaling and type conversion operations into a single kernel. By fusing these operations, the number of memory reads and writes is reduced, leading to better overall efficiency.

#### S3.3.3 Multi-threaded For Loops

We utilize multi-threading on the outer loops to distribute the workload across multiple CPU cores. Specifically, we apply multi-threading to the loops over the number of batches, number of attention heads, and number of tiles simultaneously, taking advantage of the large number of cores available in modern CPUs.

### S3.4 Using LibXSMM TPPs for accelerated modules

Our implementation uses tensor processing primitives (TPPs) [24] from the LIBXSMM library [24] to perform operator fusion and computation on tiles. TPPs use JITing to automatically generate optimized code based on underlying CPU’s instruction sets (ex. AVX, AVX2, AVX-512, AMX, etc.). Table S5 shows the set of TPPs that we use in our optimized algorithms to perform operation on tiles.

### S3.5 Supplementary Algorithms: Details of Algorithms used for accelerating Gating attention and Triangle multiplication

The Algorithms use LIBXSMM TPPs as submodules.

#### Algorithm S1

Accelerated projection kernel of gating attention for key (*K*) tensor with TPPs and tiling. While *M_I_* ∈ R*^B^*^×^*^S^* ^×^*^D^*, the algorithm is illustrated without batch dimension for simplicity.

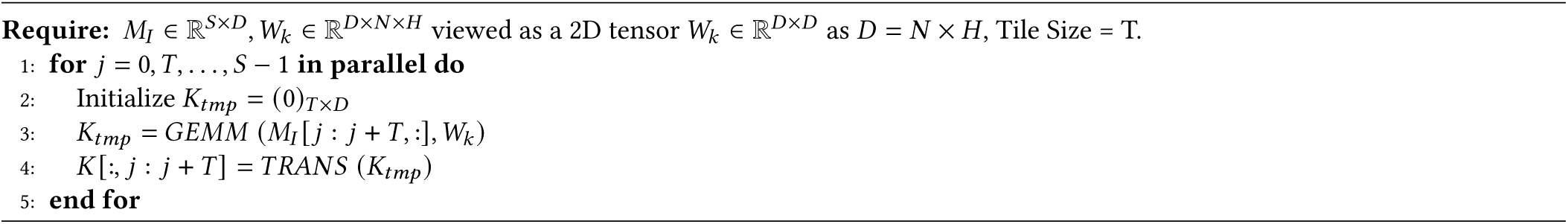

#### Algorithm S2

Accelerated SDPA kernel of gating attention with full softmax algorithm and 1D tiling. For the sake of brevity, we have skipped over the batch dimension and the scenario when the sequence length *S*is not a multiple of tile size *T_q_* and we pad the tensors to make their lengths multiple of tile size.

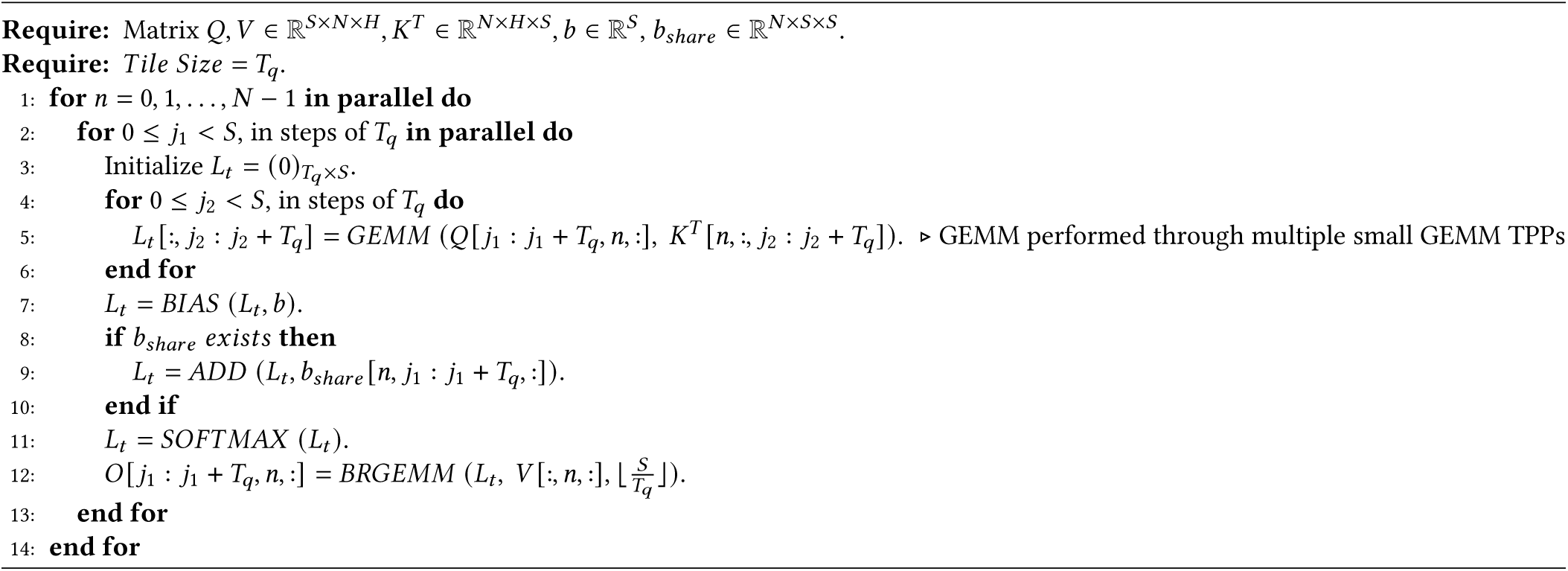

#### Algorithm S3

Accelerated SDPA kernel of gating attention with FlashAttention-2 algorithm. For the sake of brevity, we have skipped over the batch dimension and the scenario when the sequence length *S* is not a multiple of tile sizes *T, T_q_, T_kv_* and we pad the tensors to make their lengths multiple of tile size.

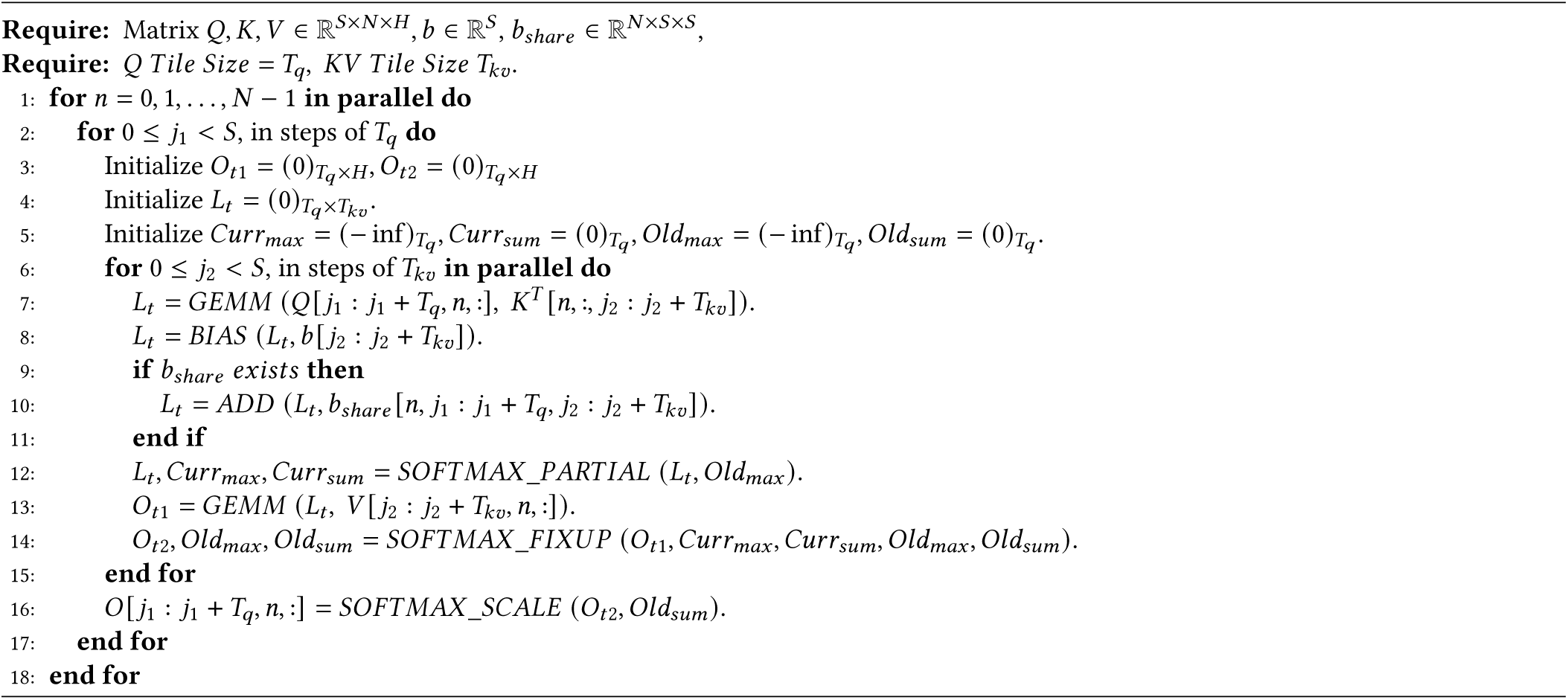

#### Algorithm S4

Accelerated Output kernel of gating attention with TPP and tiling. For the sake of brevity, we have skipped over the batch dimension.

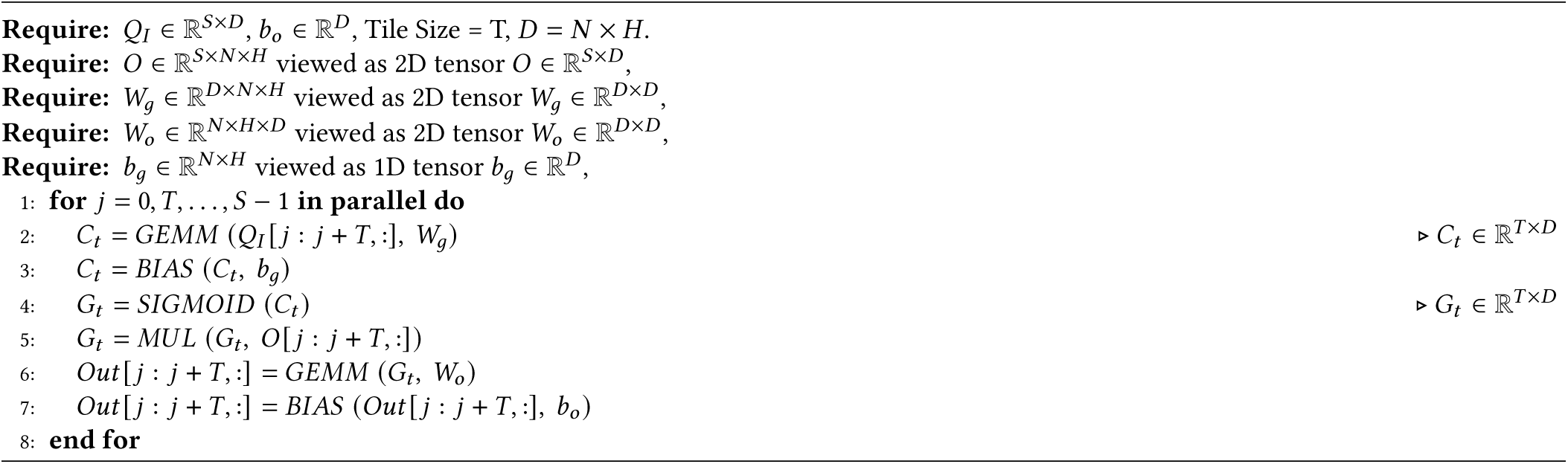

#### Algorithm S5

Accelerated projection kernel of triangle multiplication with TPPs and tiling.

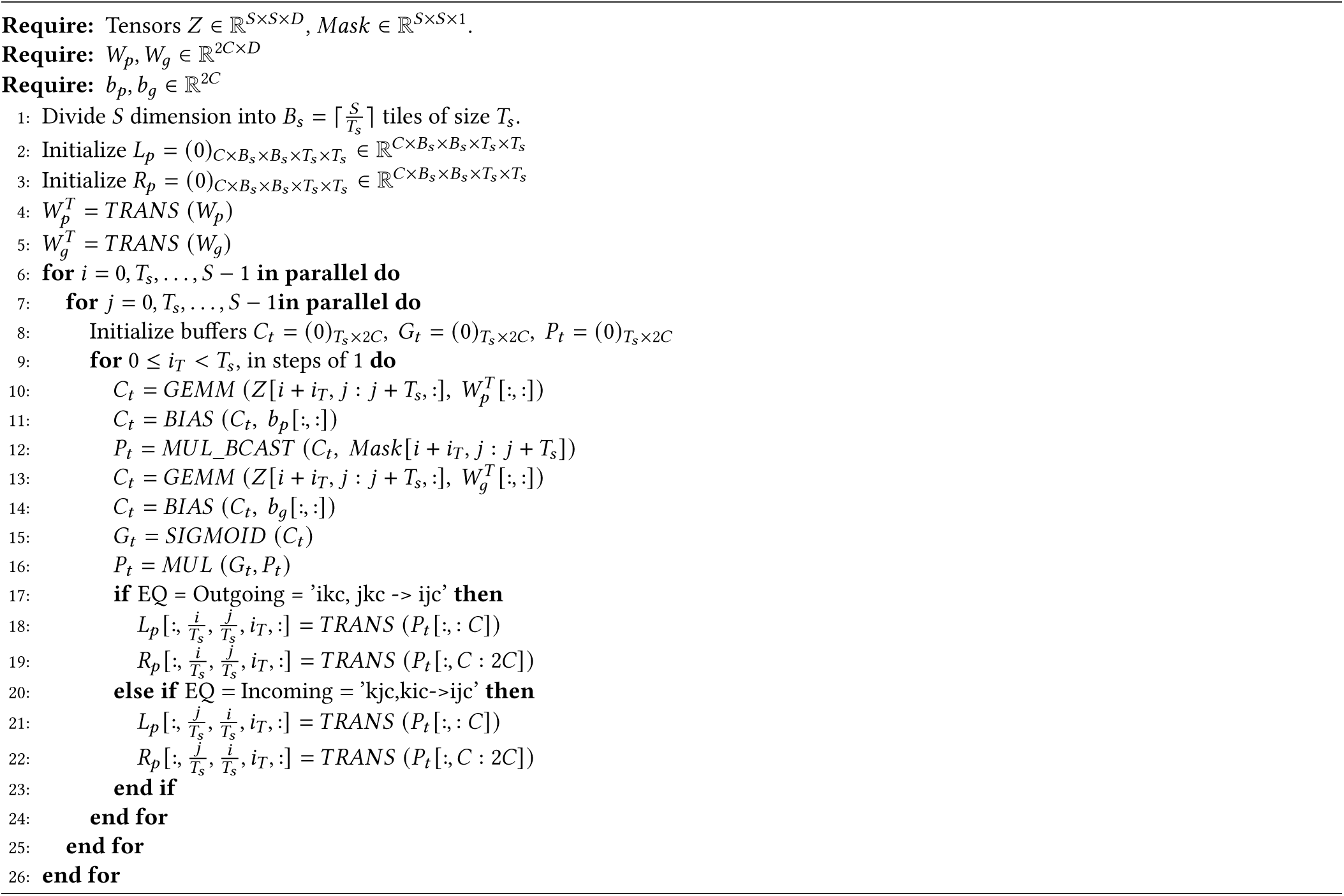

#### Algorithm S6

*Eqn_kernel* for optimized Einsum equation computation with TPPs inside the triangle multiplication module.

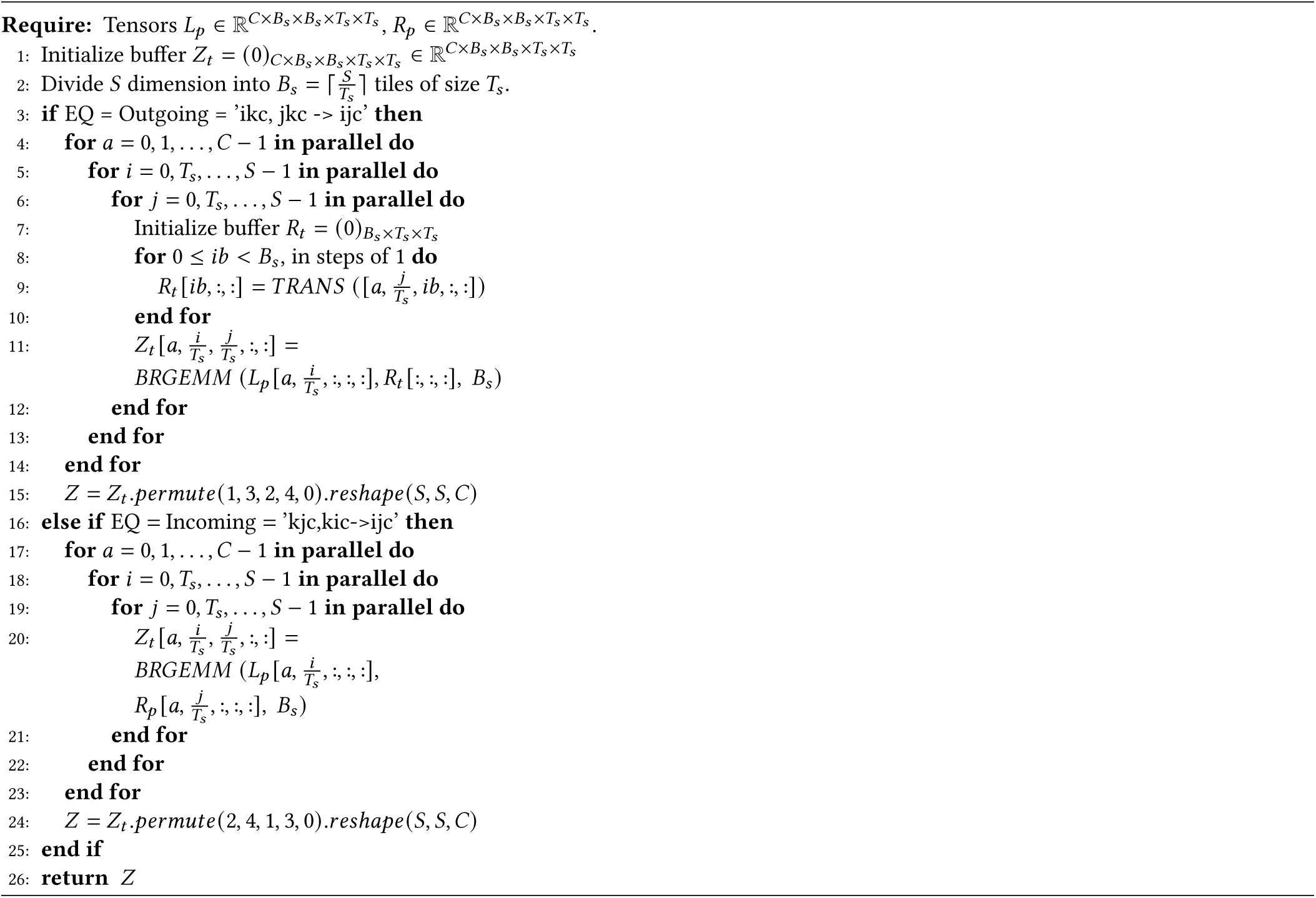

## Footnotes

1 Values cited are for matrix-multiplication without sparsity.

